# Beyond Delta Masses: MS Andrea Directly Resolves Combinatorial Peptide Modifications in Open Searches

**DOI:** 10.64898/2026.03.27.714851

**Authors:** Louise M. Buur, Stephan Winkler, Viktoria Dorfer

## Abstract

Open modification search (OMS) strategies have gained popularity in mass spectrometry-based proteomics for identification of peptides carrying unknown or unexpected post-translational modifications. However, most OMS engines report only the overall mass difference between precursor and matched peptide, without explicitly identifying or scoring combinations of modifications at the PSM level. Here, we introduce MS Andrea, a novel OMS search engine that directly identifies and scores combinations of modifications without predefining them. MS Andrea uses a sequence tag-based strategy to filter candidate peptides, evaluated using the MS Amanda scoring function. First fixed modifications only, then combinations of modifications from the Unimod database based on the observed mass shift. We evaluated MS Andrea using a human histone dataset and two phosphopeptide datasets (HeLa cells and Arabidopsis thaliana), comparing its performance with MSFragger and Sage. Across datasets, MS Andrea identified the highest number of PSMs at 1% FDR using the standard target-decoy approach while achieving higher or comparable numbers using model-based FDR estimation. Importantly, MS Andrea reports modification identities and sites for up to four modifications at the PSM level. Together, these results demonstrate that MS Andrea enables more detailed, interpretable characterization of peptide modifications while maintaining competitive identification performance in OMS-based proteomics.

## Introduction

Accurate peptide identification from mass spectrometry (MS) data is a cornerstone of modern proteomics, enabling the characterization of complex biological systems and post-translational modifications (PTMs).^1^ While conventional database search engines perform well for unmodified peptides, the growing interest in PTMs and unexpected chemical alterations has driven the development of open modification search (OMS) strategies.^2–4^ OMS approaches allow identification of peptides carrying unknown or variable modifications by tolerating large precursor mass differences, thereby expanding the detectable proteome beyond predefined modification lists.^2–4^ A prominent example of an OMS engine is MSFragger,^5^ which introduced a fragment ion indexing strategy that enables efficient database searching with wide precursor mass tolerances. This approach allows identification of peptides carrying unanticipated mass shifts without prior specification of modifications and has been widely applied for large-scale proteomics data analysis. MSFragger is supported by downstream analytical tools such as PTM-Shepherd,^6^ which aggregates mass shift observations across datasets and facilitates characterization, annotation, and localization of putative PTMs. This framework enables systematic interpretation of open search results by linking detected mass offsets to candidate modifications and their site-specific distributions. Extensions, such as localization-aware open search in MSFragger published in 2020^4^ and the very recently published “Detailed Mass Offset Search” from January 2026,^7^ index both shifted and regular ions to improve scoring and localization of unknown modifications, yet the general approach remains focused on the identification of delta masses rather than explicit multi-modification scoring. Similarly, Sage^8^ combines many of the computational innovations popularized by MSFragger with optimized performance for both narrow and wide mass tolerance searches, but it focuses on identifying a limited number of variable modifications and does not provide expanded multi-modification scoring beyond traditional delta mass identification.

Several other open search and OMS tools have been developed to improve peptide identification in the context of modifications. Open-pFind^9^ enables large-scale error-tolerant searches, identifying mass shifts above background noise. MODa^10^ and MODplus^11^ use blind, unrestrictive OMS to detect hundreds of PTMs simultaneously. TagGraph^12^ applies an unrestricted string-based search and probabilistic validation to reveal thousands of PTM types across proteomes. Despite their advances, these tools are limited in their capacity to match and robustly score more than one or two modifications per peptide, often summarizing only overall mass shifts. In parallel to open search methods, classical database search engines such as SEQUEST,^13^ Mascot,^14^ and MS Amanda,^15–17^ support variable modification handling within closed searches. These tools allow specification of some number of variable modifications per peptide, but the combinatorial explosion of candidate peptides limits the number of modifications that can be practically scored and localized. While tools like PEAKS^18,19^ incorporate de novo sequencing and PTM identification modules to capture unexpected modifications, they are often constrained by scoring models that do not scale to high numbers of simultaneous variable modifications.

Ionbot,^20^ another OMS engine, employs a machine-learning–based scoring model and supports searches with up to two specified variable modifications, in combination with one additional unexpected modification drawn from the Unimod^21^ database. This hybrid strategy allows identification of peptides carrying modifications that were not anticipated in the presence of expected ones. However, the total number of modifications that can be explicitly modeled and scored per peptide remains limited to two or three, restricting sensitivity for peptides carrying more complex modifications. The number of modifications that co-occur on a single peptide is, however, also limited, even for the most heavily modified proteins. For histone N-tails, a classic example of proteins with combinatorial modifications, individual bottom-up peptides usually have two or three, but sometimes more, modified residues, despite the N-tails collectively carrying more than that.^22^ Hyperphosphorylated proteins follow the same pattern. The Tau protein is phosphorylated at more than 50 residues in disease states,^23^ but the modifications are distributed across the sequence, so only a few are found on any single peptide. Capturing such cases therefore requires scoring more than the two or three modifications that current search engines can identify, but rarely more than four in bottom-up experiments.^24,25^ Allowing modifications beyond this combinatorially expands the search space without necessarily capturing additional biology, lowering sensitivity and increasing the false discovery rate (FDR).^26,27^ Here we present MS Andrea, a new OMS engine that enables matching and scoring of up to four distinct variable modifications per peptide without having to explicitly specify them and directly reports PTMs at the peptide spectrum match (PSM).To address the large search space inherent to OMS, MS Andrea employs a sequence tag–based strategy to efficiently reduce the number of candidate peptides subjected to comprehensive scoring.

## Methods

### Overview of the MS Andrea search algorithm

An overview of the MS Andrea search algorithm is shown in Figure 1. First, each spectrum is pre-processed, which includes the removal of the precursor peaks and simple deconvolution (deisotoping and charge reduction), followed by precursor mass-dependent peak picking for finding sequence tags. Separately, peaks are also picked for scoring in the same way as they are in MS Amanda.^15^ Sequence tags of two, three and four amino acid (AA) residues are then extracted from each spectrum. Sequence tag finding is described in detail in the following section. Peptide candidates from the database are then going through a two stepped filtering, first based on the sequence tags and then score. All peptides that contain at least one sequence tag of three or four residues or two sequence tags of two residues are selected for further filtering using a wide MS1 tolerance to accommodate for the mass of potential variable modifications of the peptide. The remaining peptide candidates are then filtered based on score, where each of the peptide candidates is scored using the MS Amanda^15^ scoring function, considering only fixed modifications specified in the search settings. The top ten scoring PSMs are then considered in the main scoring. Each peptide candidate will be scored with variable modifications that have a combined mass corresponding to the delta mass between the precursor and the matched peptide from score-based filtering. MS Andrea will match and score combinations of up to four modifications from the Unimod ^21^ database per peptide. For peptides carrying no or unknown modifications, the unmodified peptidoform will be reported alongside a potential delta mass. All spectra are searched against both the target and a decoy database of reversed peptides, re-ranked according to score, and exported to a tab-separated .csv file.

**Figure 1:**
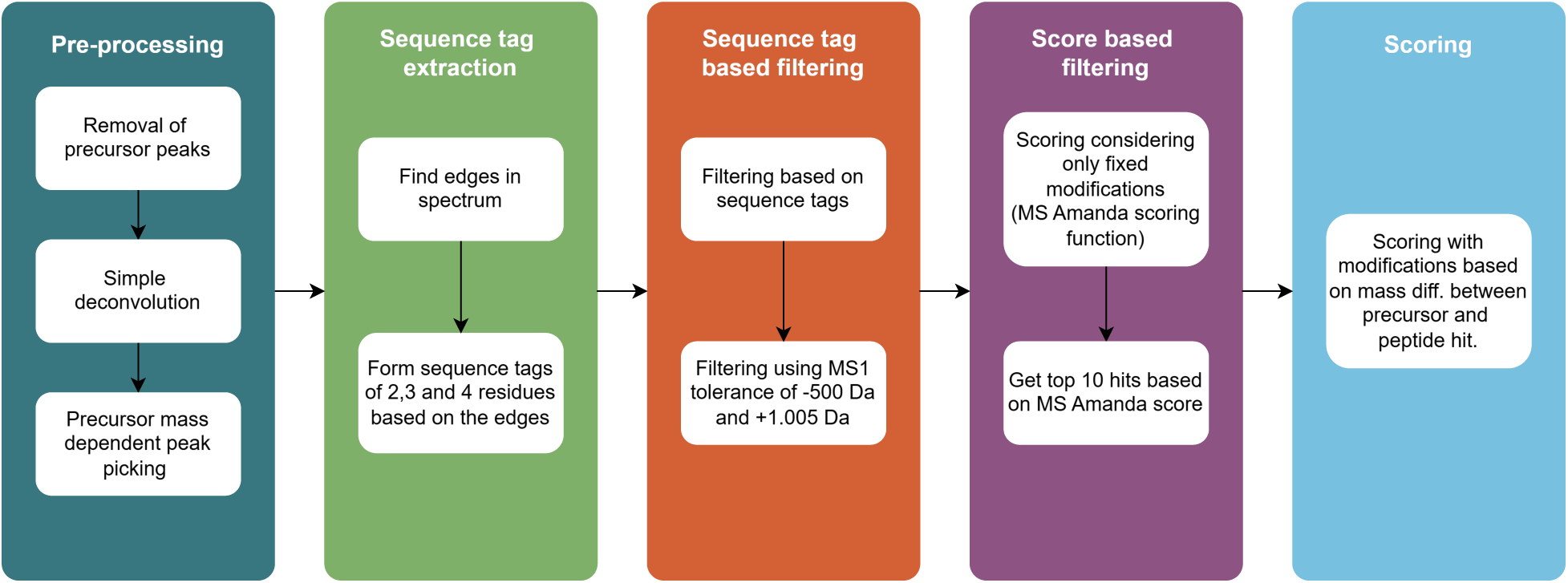
Overview of the MS Andrea peptide identification workflow. MS/MS spectra are initially pre-processed by removing precursor ion peaks, performing deconvolution, and selecting fragment peaks based on precursor mass. Subsequently, edges are identified between peaks whose delta mass corresponds to amino acid residues, enabling the generation of sequence tags of length two, three, and four residues. These sequence tags are used to filter peptide candidates, followed by additional filtering based on an MS1 precursor mass tolerance window of -500 Da and +1.005 Da. The remaining candidates are further filtered by retaining the top 10 candidates after being scored using the MS Amanda^15^ scoring function considering only fixed modifications. In the final stage, these candidates are scored again with consideration of potential modifications corresponding to the mass between the precursor and the matched peptide from the filtering round, resulting in the final ranked peptide identifications.

#### Simple deconvolution

After the removal of the precursor peaks, the spectrum undergoes simple deconvolution in two steps: deisotoping and charge reduction. In the deisotoping step, MS Andrea will search for peaks in the spectrum that correspond to isotopic distributions of peaks with a charge up to 3+. If such isotopic distributions are found, they will be reduced to the monoisotopic mass of the isotope with the summed intensities. In addition, intensities of doubly/triply charged peaks will be added to the singly charged one, and the doubly/triply charged peak will be removed from the spectrum. If the singly charged peak is not present, the m/z of the singly charged species will be artificially added to the spectrum.

#### Precursor mass-dependent peak picking for sequence tag identification

Following deconvolution, peaks are picked from each spectrum based on the uncharged precursor mass, which are then used for finding sequence tags. The peak picking is defined as:

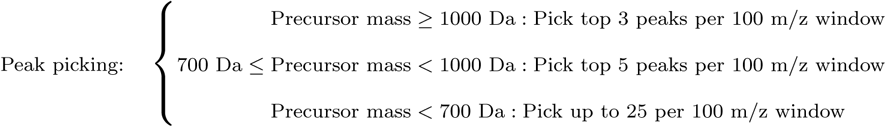

For spectra with an uncharged precursor mass of more than 1000 Da, the top 3 most intense peaks per 100 m/z window are picked. If the uncharged precursor mass is more than 700 Da but less than 1000 Da, the top 5 most intense peaks will be picked from each 100 m/z window. For spectra with an uncharged precursor mass less than 700 Da, all or up to 25 peaks are picked from each 100 m/z window. This precursor mass-dependent peak picking was empirically optimized to balance information content and noise reduction. For higher-mass peptides (*≥* 1000 Da), limiting selection to the three most intense peaks preserves the most informative fragment ions and is sufficient for reliable sequence tag generation. In contrast, for lower-mass peptides, particularly those with an uncharged precursor mass *<*700 Da, more information and, therefore, more peaks are needed to find reliable sequence tags.

After peak picking, auxiliary peaks, serving as the start and end points of the fragment ion series, are added to the list of selected peaks according to the fragmentation method specified in the settings file. For instance, if b-and y-ions are the specified ion types in the settings file, four auxiliary masses: (1) *m*_proton_, (2) *m*_H_2_O_ + *m*_proton_, (3) *m*_precursor_ + *m*_proton_ and (4) (*m*_precursor_ + *m*_proton_) - *m*_H_2_O_, are added to the list of selected peaks that are used when finding sequence tags. See Supplementary Table S1 for a full list of added auxiliary peaks.

#### Sequence tag extraction

The sequence tag extraction implemented in MS Andrea is derived from the principle of the spectrum graph approach traditionally used in de novo peptide sequencing, first described by Bartels.^28^ In MS Andrea a set of connections between peaks in the spectrum that are displaced by the masses of the 20 canonical AAs, which we will refer to as “edges”, is constructed, as shown in Figure 2A and 2B.

**Figure 2:**
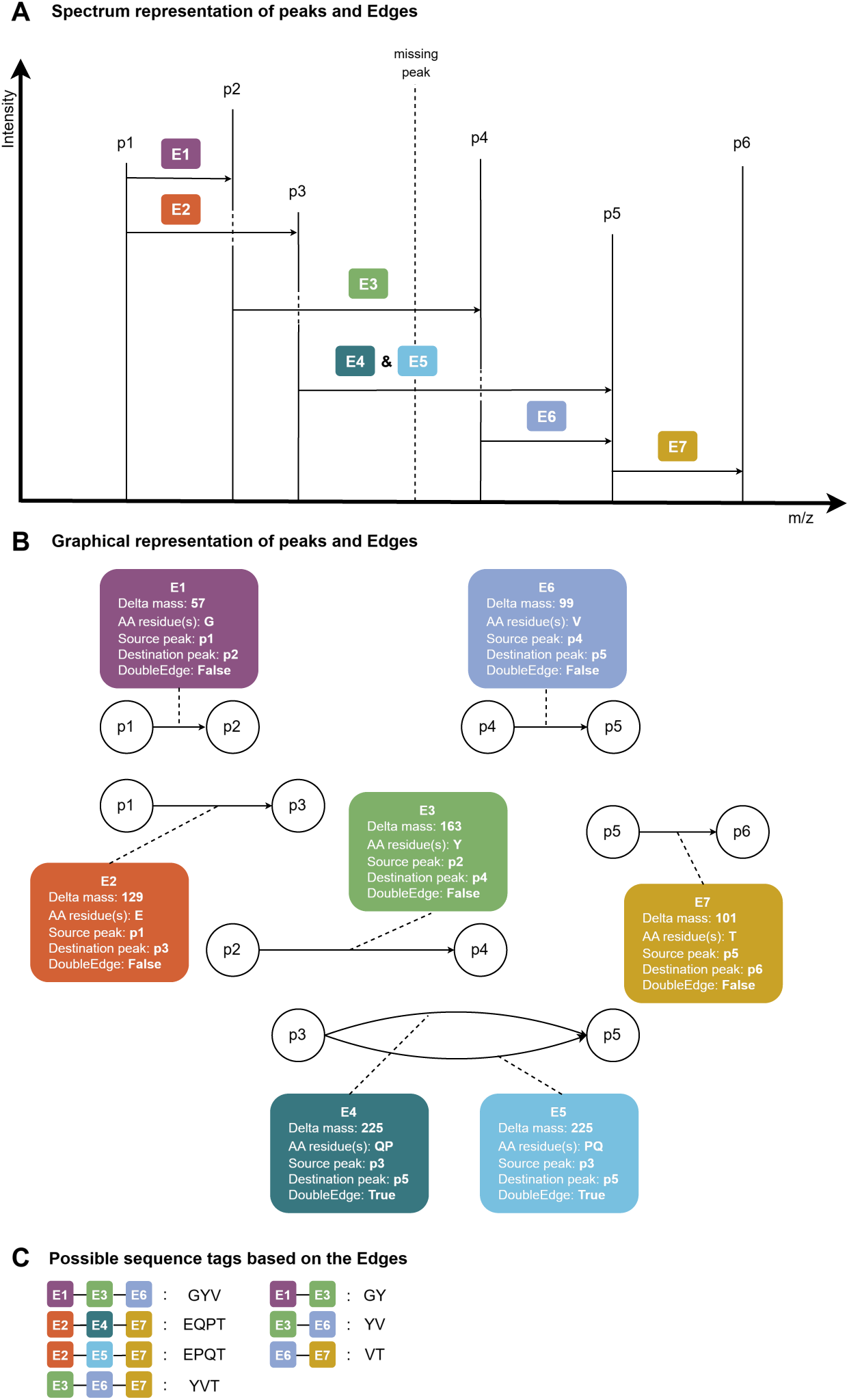
Spectrum and graphical representation of edges and their role in forming sequence tags in MS Andrea. (A) Spectrum representation showing connections, which we refer to as edges, between peaks whose delta masses correspond to one or two amino acid (AA) residues. Shown here are seven edges (E1-E7) between six peaks (p1-p6). (B) Graphical representation of the edges. Each edge is defined by the delta mass between a source peak, a destination peak and the corresponding AA residue(s). If the delta mass corresponds to the combined mass of two AA residues, the “DoubleEdge” attribute is set to True (e.g., E4 and E5). If the delta mass corresponds to a single AA residue, the “DoubleEdge” attribute is set to False. (C) Sequence tags that can be generated based on the edges shown in panels A and B. Theoretically, two additional double edges could be formed, namely between p2 and p5 as well as between p4 and p6. However, they are not shown in the figure for the sake of simplicity.

Each edge is defined by a delta mass (dm) corresponding to the mass of one or two AA residue(s), a source (src) peak, a destination (dest) peak, the AA residue name(s) corresponding to the dm, and lastly, whether or not the edge is a double edge (has an dm corresponding to two AA residues). For double edges, it is unknown which AA residue comes first in the sequence. Thereby, if a double edge has an dm corresponding to the combined mass of, for instance, proline and glutamine, the AA residue attribute is set to “PQ” for this edge. Then a second edge is created with the exact same attributes as the first one, but with the AA residue attribute set to “QP” instead, as illustrated for E4 and E5 in Figure 2B. Mathematically, an edge between two peaks, p1 and p2 is defined as *e* := *edge*(*p*1*, p*2), and its validity is defined as follows:

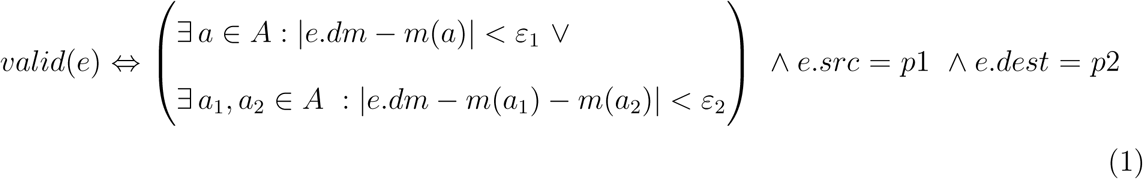

where *ε*_1_ is the mass tolerance for single edges, *ε*_2_ is the mass tolerance for double edges, *A* is the set of the 20 canonical AA residues and *a* is one of these AA residues. This means that an edge is valid if it connects two peaks observed in the spectrum and the delta mass between those peaks can be explained by either one or two AA residues (within small tolerances *ε*_1_ and *ε*_2_, respectively). *ε*_1_ and *ε*_2_ are by default set to 0.015 Da and 0.004 Da, respectively. These tolerances were empirically optimized based on Orbitrap datasets to obtain as many correct sequence tags as possible while keeping false ones at a minimum. These tolerances can be changed by the user in the settings file. For data acquired on instruments with time-of-flight (TOF) mass analyzers (e.g., Bruker timsTOF or SCIEX ZenoTOF) and for data acquired on a Thermo Fisher Scientific Orbitrap Astral Mass Spectrometers, we advise using slightly wider tolerances than the Orbitrap defaults (e.g. 20-40 ppm for TOF and 20 ppm for Astral data, depending on the acquisition settings).

Based on the edges, sequence tags of two, three and four AA residues are then constructed, as shown in Figure 2C. A sequence tag is defined as *tag* := {*e_i_*}, |*tag*| ∈ {2, 3} and its validity is defined as:

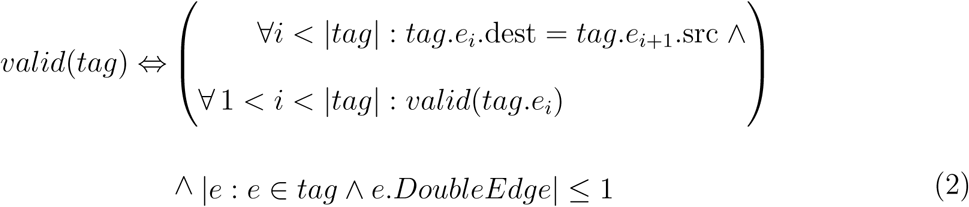

Valid sequence tags of MS Andrea are then based on either two of three edges, corresponding to three or four consecutive peaks in the spectrum. Only one edge can be a double edge, given that the sequence tag is based on three edges. Sequence tags that are based on three edges can therefore be presented by three or four AA residues. Sequence tags of length two would be possible to be formed based on a single double edge, but we concluded that only two consecutive peaks in the spectrum, especially peaks displaced by the mass of two AA residues, do not provide enough evidence for a reliable sequence tag. Similarly, the higher mass ambiguity that is associated with the double edges is the reason that not more than one double edge is allowed per sequence tag. We chose to include double edges to accommodate cases in which a peak is missing that would otherwise be a potential dest and src peak of two adjacent single edges, e.g., a low-intensity peak that was not retained during peak picking or a peak that is simply not present in the spectrum. Such a case is illustrated in Figure 2A where the dashed peak indicates a missing peak, where a double edge is formed between p3 and p5 to compensate for the missingness.

Sticking to single edges only could in worst case scenarios lead to situations, where no sequence tags can be formed for the spectrum, particularly for short peptides. This means that no peptide candidates would be found and the spectrum would be skipped, potentially meaning a missed identification.

To further ensure that “correct” sequence tags can be found, the fixed modifications that are specified for the search are also considered in the tag finding for the affected residues. For instance, if carbamidomethylation of cysteines is defined in the settings as a fixed modification, the mass of cysteine will be replaced by the mass of cysteine plus the mass of carbamidomethyl. Due to its high prevalence, oxidation of methionine is considered as a variable modification for tag finding per default. Edges labeled with the AA residue “M” are therefore created when the dm between the src and dest peaks is equal to both methionine and the mass of methionine plus an oxidation. We found that including this modification increases the number of sequence tags that match the sequence determined by a closed search with MS Amanda (data not shown). Because leucine and isoleucine are indistinguishable by mass, two edges will be created for peaks that are displaced by the mass of leucine and isoleucine. One edge will have “L” as the AA residue attribute and the other “I”. The remaining attributes will be the same for the two edges. This ensures that both options of “L” and “I” are taken into account when the sequence tags are used for filtering (see next section).

#### Sequence tag based filtering

After finding sequence tags, MS Andrea will use these to filter the potential candidates in the peptide database. It will collect all peptide candidates that contain at least one sequence tag of three or four residues or at least two sequence tags of two residues, like it is shown in Figure 3. As it is unknown if a sequence tag was found based on a series of a/b/c-or x/y/z-ions, target and decoy peptide candidates that contain the original tag sequence or the reversed tag sequence will be collected. For example, if the tag is “TAG”, sequences containing either “TAG” or its reverse (“GAT”) are retained for the next filtering step.

**Figure 3:**
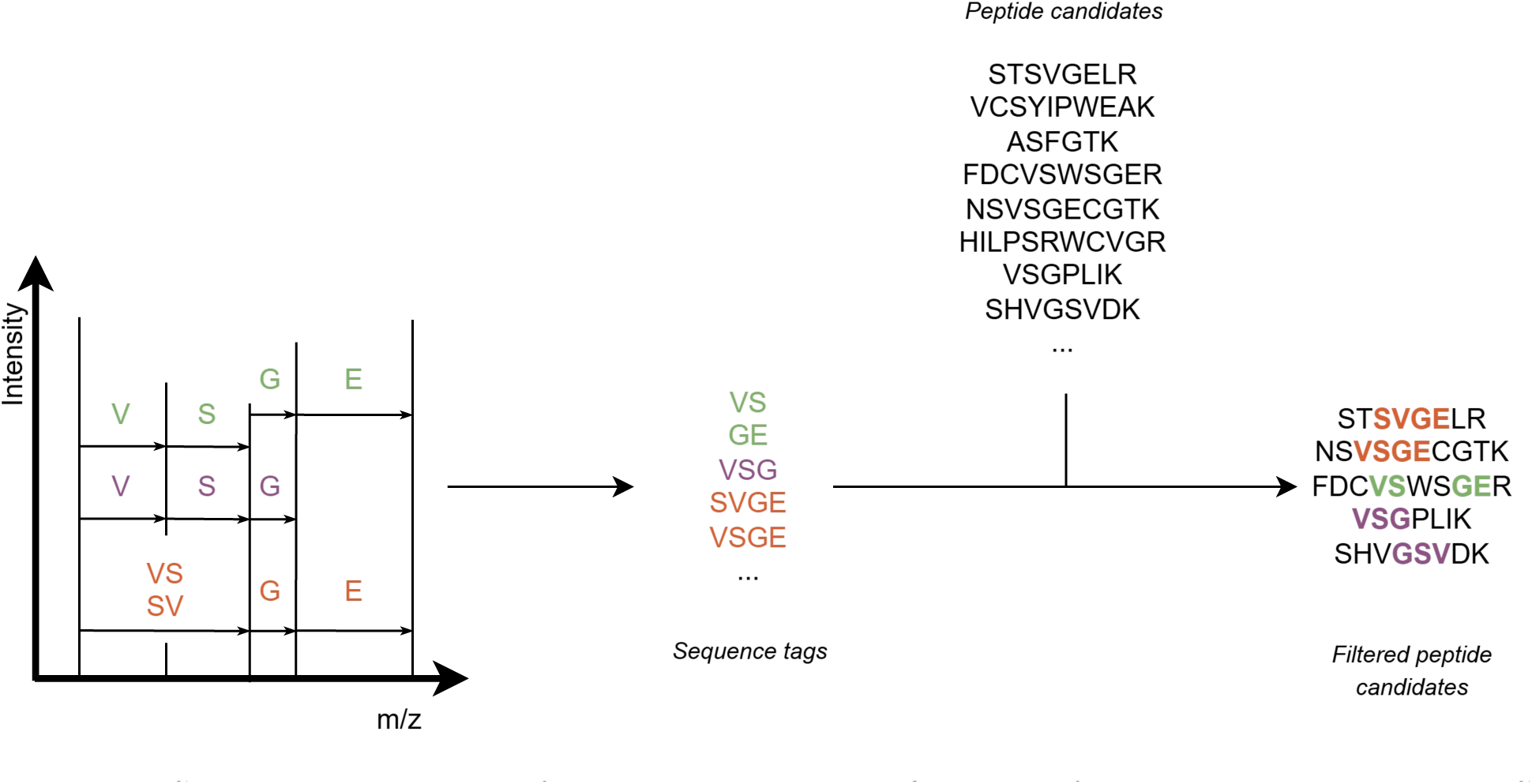
Schematic overview of sequence tag based filtering of peptide candidates in MS Andrea. Peaks in the spectrum (left) that are displaced by the masses of amino acids are connected to form sequence tags: VS, GE, VSG, SVGE and VSGE. These sequence tags are then matched against the list of peptide candidates from the database (middle). Peptide candidates that contain at least one sequence tag of three or four residues and peptide candidates that contain at least two sequence tags of two residues are retained for further filtering. Because it is unknown whether a sequence tag is formed based on edges between peaks that represent b-or y-ions in the spectrum, peptide candidates that contain the reversed sequence tags are also retained, as shown in the last row of the filtered peptide candidates (right).

Next, the remaining peptide candidates will be filtered using a wide MS1 tolerance of -500 Da and +1.005 Da to enclose a variety of combined modifications present in the Unimod database.^21^ These tolerance values are set as defaults but can be modified by the user via the settings file. The upper MS1 tolerance limit of +1.005 Da was selected to account for isotopic mass misassignments occasionally introduced by the MS instruments, as the most common misassignment arises from selection of the ^13^C isotopic peak rather than the ^12^C, which introduces a mass shift of approximately +1.003355 Da.

#### Score Based Filtering

Scoring in MS Andrea is performed using the MS Amanda^15^ scoring function *S*(*s, pep*), as defined below.

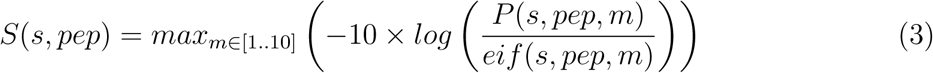

where, *P* (*s, pep, m*) reflects the reliability of a PSM under the null hypothesis that the match occurred randomly, assuming a binomial distribution, and *eif* (*s, pep, m*) reflects the explained ion current. Please refer to the original MS Amanda publication for details. First, each peptide candidate (after previous filtering steps) is evaluated against the spectrum, considering only fixed modifications. Subsequently, the search results are filtered so that only the top ten highest-scoring peptides remain (score-based filtering step in Figure 1). Keeping the top ten is the default value; this can be changed by the user in the settings file.

#### Construction of Modification Dictionary

Filtered peptides are then re-evaluated in the main scoring step with variable modifications. First, a dictionary of possible combinations of up to four modification types is generated once based on the Unimod file.^21^

As of June 2026, the Unimod database contains 1358 entries. Considering all four-modification combinations would therefore require evaluating *C*(1358, 4) *≈* 1.41 *·* 10^11^ combinations, which is not computationally feasible within a reasonable amount of time. To avoid this combinatorial explosion, MS Andrea applies four filters before the modification dictionary is built. This filtering process is described in Supplementary Element S1. Together, these filters reduce the pool of modifications from 1358 to 133 commonly observed, relevant modifications, and the number of four-modification combinations drops from *∼* 1.41 *·* 10^11^ to *C*(133, 4) = 12, 475, 445, which is roughly four orders of magnitude smaller.

This number is then further reduced to 1,902,572 combinations using the default open search precursor mass tolerances (-500 Da and +1.005 Da) and removing combinations with masses that cancel out each other.

A full list of the 133 modifications considered by MS Andrea by default (modifications that have at least one residue classified as “Post-translational”) is provided in Supplementary Table S2. Optionally, users can expand this set to also include modifications with residues classified as “Chemical derivative” and/or “Artefact”using the considered unimod classifications in the settings file. Including additional classifications will, however, result in a larger number of considered combined modifications and therefore a longer search time.

#### Scoring

For each peptide *pep*, the set of modifications *mod*(*pep,* Δ) that is evaluated is defined such that the combined mass matches the delta mass between the uncharged precursor and the peptide identified in the first round. For each combination of modifications, *m* in *mod*(*pep,* Δ), the peptide, the modification combination is calculated as *pep_mod_m__*. Then, the final score *S*_2_(*pep*), is determined by evaluating each peptide with each modification combination against the spectrum and scored using the MS Amanda scoring function:

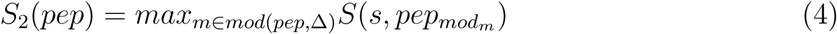

Because the empty modification set is always included in *mod*(*pep,* Δ) Equation 4 evaluates the unmodified peptide alongside all modification sets whose combined mass matches the observed mass shift and reports whichever maximizes the score. The unmodified peptide is therefore reported along with the observed mass shift whenever no candidate modification set outscores it. This is the case for peptides carrying modifications that are not represented in the Unimod database or when the unmodified sequence is simply the best match to the spectrum. MS Andrea does not attempt to localize unknown mass shifts.

### Benchmark Datasets and Search Parameters

To assess the performance of MS Andrea, we compared the search results from MS Andrea to those of MS Fragger and Sage, using three different data sets from different species to demonstrate the applicability of MS Andrea. We analyzed the different datasets using MS Andrea v.1.0, MSFragger v.4.4.1 within the Fragpipe platform v.24.0 with PeptideProphet^29^ and PTM-Shepherd^6^ v.3.0.11, and Sage v0.14.7.

In addition, one dataset of synthetic phosphorylated peptides was analysed with MS Andrea only to evaluate how MS Andrea controls FDR. For entrapment analysis, FDRBench^30^ v. 1.1.1 was used to add shuffled entrapment sequences to the FASTA file used for the search. The datasets described in the following sections were used for benchmarking MS Andrea and not for testing and development. For that purpose, two independent datasets were used. (1) Peptides from HeLa cells published by Dorfer et al. 2018,^31^ available via the PRIDE repository^32^ (https://www.ebi.ac.uk/pride/) using identifier PXD007750. (2) Phosphorylation-enriched peptides from HT29 cells, published by Piersma et al. 2015,^33^ available with identifier PXD001550 via the PRIDE repository.

#### Phosphorylated peptides derived from HeLa cells

In 2020, Bekker-Jensen et al. published their work on phosphoproteome profiling using direct data-independent acquisition and compared their approach to a standard data-dependent acquisition (DDA) approach.^34^ From this dataset, six samples from the LFQphos LCMS data were downloaded and analyzed with the three search engines. See Supplementary Element S2 for the full list of the analyzed files. The complete dataset is available via the PRIDE repository under identifier PXD014525. The human proteome (UP000005640) was downloaded from UniProtKB^35^ as a FASTA file on 04.11.2025 and was used for the searches. Additionally, the second replicate of this dataset was used for a search where phosphorylation was allowed on all 20 amino acid residues to evaluate the localization accurary of MS Andrea.

#### Phosphorylated peptides derived from *Arabidopsis thaliana*

Mergner and colleagues published a comprehensive atlas of the *Arabidopsis thaliana* proteome in 2020.^36^ Eight raw files with enriched phosphopeptides from the flower of *Arabidopsis thaliana* were downloaded and analyzed with the three search engines. See Supplementary Element S3 for the full list of the analyzed files. The full dataset is available on the PRIDE repository with identifier PXD013868. For the searches, the FASTA file provided by the authors was used.

#### Histones from Human Breast Cancer Tissue

Noberini et al. published their method, PAThology tissue analysis of Histones by Mass Spectrometry (PAT-H-MS) in 2016. They tested their method on histone proteins derived from tissue from breast cancer patients.^37^ From this dataset, three raw files were downloaded and analyzed. See Supplementary Element S4 for the full list of the analyzed files. The complete dataset was made available by the authors on the PRIDE repository under project identifier PXD002669. For the searches, a FASTA file containing all human H3 and H4 histones was used.

#### Synthetic Phosphorylated Peptide Library

Marx et al. 2013^38^ published a library of synthetic unmodified and phosphorylated peptides. They constructed the library based on tryptic phosphopeptides observed in five large-scale studies of phosphorylations in human samples. The full library consists of 96 different seed peptides that served as a permutation template to create peptides with permutated phosphosites (serine, threonine and tyrosine) and the flanking residues (all 20 canonical AAs). Three libraries were randomly chosen and analysed with MS Andrea only. See Supplementary Element S5 for the full list of the analyzed files. The full library is available on the PRIDE repository using project identifier PXD000138. The FASTA file provided in the repository was used for the searches.

### Search Parameters

All raw files were converted to .mzml format using ThermoRawFileParser^39^ v1.4.5 prior to the database searches. For a fair comparison, the search parameters were set to be the same or as similar as possible for all three search engines.

A tryptic digest was applied to both the HeLa and *Arabidopsis thaliana* datasets; therefore, trypsin was specified as the enzyme with up to two missed cleavages (four missed cleavages were specified for the synthetic peptide library). For the histone dataset, Arg-C digestion was specified. Minimum peptide length was set to 7, and maximum peptide length was set to 30. b-and y-ions were selected as the ion types. The lower limit open search precursor mass tolerance was set to 500 Da for the HeLa and *Arabidopsis thaliana* datasets, for the histone dataset, it was set to 200 Da and to 100 Da for the synthetic peptide library dataset. The upper mass tolerance was set to 1.005 Da for all datasets. The fragment ion tolerance was set to 0.02 Da for the HeLa, *Arabidopsis thaliana* and the synthetic peptide library datasets and to 6 ppm for the histone dataset (which was the tolerance used by Noberini et al. in their analysis). Carbamidomethyl (+57.02146 Da) of cysteine residues was set as a fixed modification for the HeLa and *Arabidopsis thaliana* datasets. No fixed modifications were set for the histone dataset and the synthetic peptide library dataset. The maximum number of modifications per peptide was set to four. Deisotoping was enabled for all three search engines. Removal of the precursor peak was enabled in MS Andrea and MSFragger (not a parameter for Sage). Common contaminants from the cRAP^40^ database were added to the FASTA files before processing with MS Andrea and Sage. For searches with MSFragger, decoy sequences and common contaminants were added to the FASTA files intrinsically in FragPipe v. 24.0.

Post-processing of the search results was done using custom R-and Python scripts and adjusted to 1% FDR using the standard target-decoy approach (STDA) for all three search engines. For STDA, the MS Amanda score was used for MS Andrea and the Hyperscore was used for MSFragger and Sage. For model-based FDR estimation, Percolator^41^ v. 3.06.1 was used for MS Andrea results. Pin files were generated from MS Andrea output files with a custom Python script. For MSFragger, q-values assigned by PeptideProphet were utilized, whereas the spectrum level q-value assigned by Sage itself were used for Sage results.

## Results

### MS Andrea identifies high numbers of PSMs

We here present MS Andrea, an OMS engine that scores and identifies up to four variable modifications per peptide and reports the modification types and sites directly on the PSM level. To evaluate the performance of MS Andrea, we tested it on three different datasets and compared the search results with those of MSFragger and Sage, two well-known and widely used OMS engines. The average number of PSMs identified for the three replicates in the Histone dataset is shown in Figure 4. Search results were adjusted to 1% FDR using both the standard target-decoy approach (STDA), shown with orange bars, and model-based post-processing either with Percolator^41^, PeptideProphet^29^ or intrinsically in Sage, which is shown in green bars. In both cases, MS Andrea achieved the highest number of PSMs, with an average of *∼*540 PSMs for the three replicates using the STDA for FDR adjusting and 1000 PSMs per replicate at 1% FDR when combining MS Andrea with Percolator. MSFragger found on average 418 PSMs per replicate in combination with PeptideProphet and 285 PSMs on average at 1% FDR using STDA. In contrast, Sage identifies a very low number of PSMs of 24 on average using STDA. However, when using the spectrum level q-value assigned by Sage itself for FDR adjusting the number of identifications drastically increased to 677 on average. See Supplementary Table S3 for a full overview of the exact numbers. For the HeLa and *Arabidopsis thaliana* datasets, a similar pattern can be observed when comparing the number of identifications at 1% FDR based on STDA. MS Andrea finds the most PSMs for both datasets and Sage the fewest (Supplementary Figure S1 and S2 and Supplementary Table S4 and S5). Generally, the number of identifications by Sage using STDA is on the low side compared to MS Andrea and MSFragger. This observation is what initially led us to evaluate the search results using model-based post-processing. Originally, we wanted to compare the search engines purely based on their individual performance using STDA for FDR adjustment. However, we figured that it would be the most fair to utilize the q-values assigned by Sage and then compare the number of identifications based on q-values assigned by Percolator for MS Andrea and PeptideProphet for MSFragger. Comparing the number of identifications at 1% FDR using model-based post-processing, all three search engines achieve comparable numbers of identifications for the *Arabidopsis thaliana* dataset, although with slightly more identifications by MS Fragger with PeptideProphet (Supplementary Figure S1 and Supplementary Table S4). For the HeLa dataset MSFragger with PeptideProphet also acheives the highest number of identification, followed by MS Andrea with Percolator and then Sage with spectrum level q-values (Supplementary Figure S2 and Supplementary Table S5).

**Figure 4:**
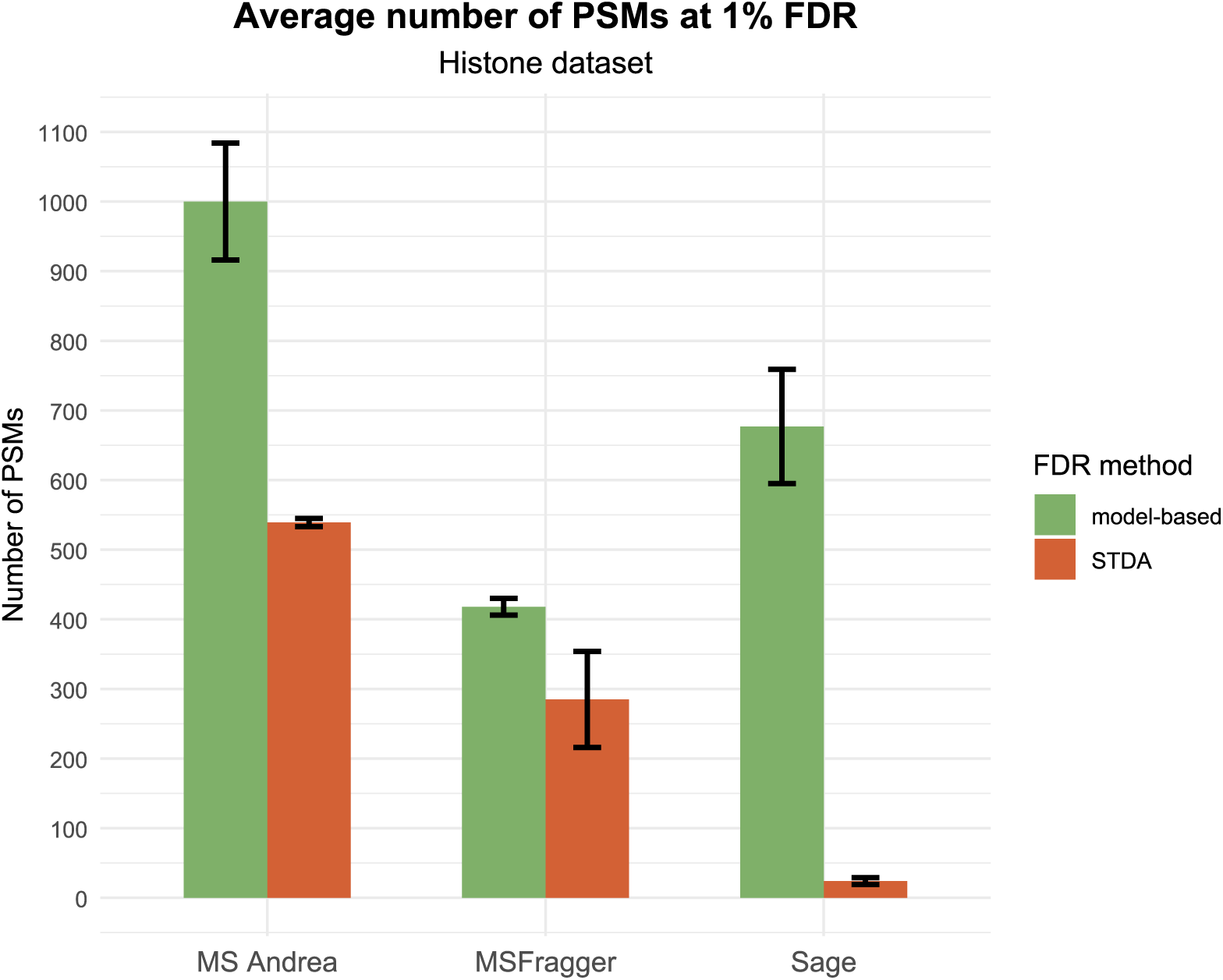
Average number of PSMs determined by MS Andrea, MSFragger and Sage for the three replicates of the Histone dataset. Green bars show the average number of PSMs for the search engine with model-based post-processing. For MS Andrea post-processing was done with Percolator,^41^ and for MSFragger, PeptideProphet^29^ was used within Fragpipe For Sage results, the q-value determined on spectrum level by Sage was used. Orange bars show the average number of PSMs at 1% false discovery rate (FDR) using the standard target-decoy approach (STDA). Error bars show standard deviation among the replicates.

### Overlap in identified Peptide Spectrum Matches

We compared the overlap of PSMs identified in the first replicate of the Histone dataset at 1% FDR after model-based post-processing, as shown in Figure 5. The Venn diagram shows that there is a 9.6% overlap, corresponding to 156 PSMs, between the three search engines. MS Andrea identified the highest number of unique PSMs, with nearly 700, followed by Sage with 361 and MSFragger with just under 100. Given the lower number of identifications by MSFragger, it also shows the lowest overlap with the other search engines, sharing 4% of its PSMs with MS Andrea and 5.3% with Sage. In contrast, the overlap between Sage and MS Andrea is 10.4%, approximately twice as high.

**Figure 5:**
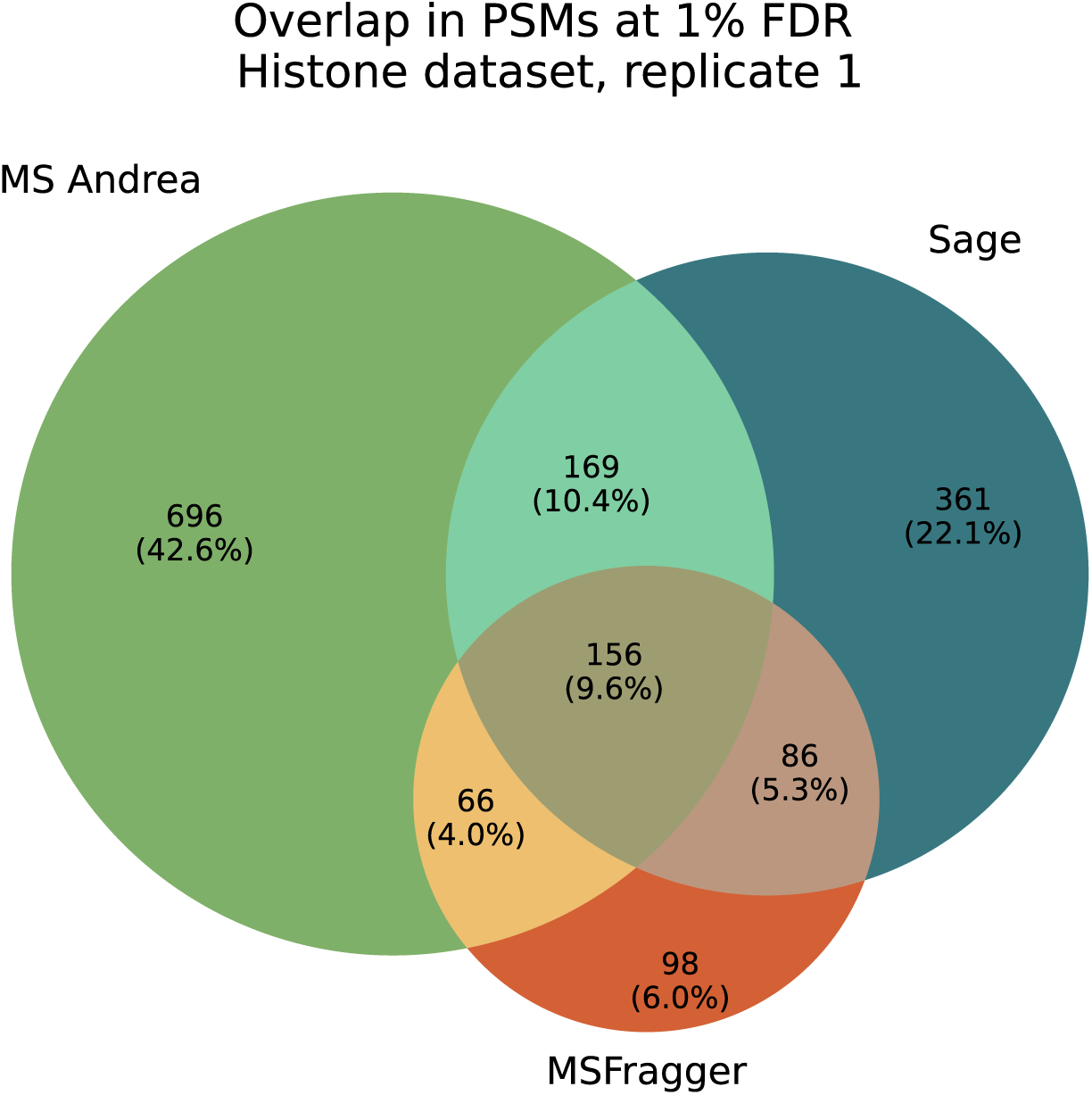
Overlap in peptide spectrum matches (PSMs) identified by MS Andrea (green), MSFragger (orange) and Sage (dark blue) at 1% false discovery rate (FDR) for the first replicate of the Histone dataset. The overlap is based on peptide-level q-values determined by Percolator (for MS Andrea), PeptideProphet (for MSFragger) and Sage, respectively.

Overall, these results indicate that, despite differences in the PSMs identified by each search engine, there is a considerable degree of agreement between them. While the large number of PSMs uniquely identified by MS Andrea and Sage is notable, further investigation revealed that some of the discrepancies arise because one search engine assigns a spectrum to a target hit whereas the other assigns it to a decoy hit. This suggests that at least part of the observed disagreement is due to differences in target–decoy classification rather than entirely distinct spectrum assignments.

### MS Andrea identifies modifications on PSM level

The strength and advantage of MS Andrea is its capability of identifying modifications on the PSM level. Due to how the MS Andrea algorithm is designed, it will, based on the delta mass between the precursor and the peptide from the database, match and score all possible combinations of up to four modifications that have a combined mass that corresponds to the delta mass. An example of this is shown in Table 1.

**Table 1:**
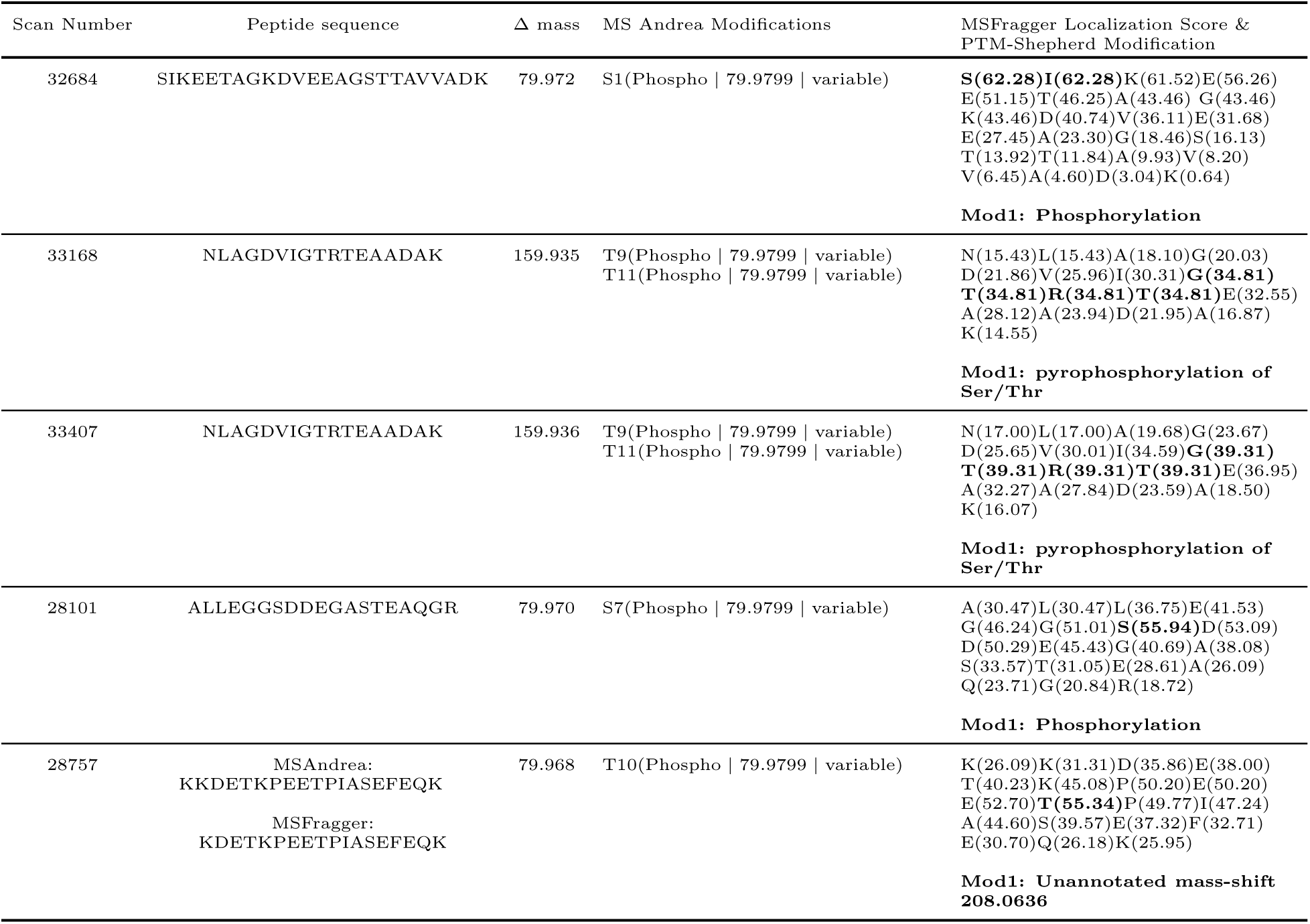
The five highest-scoring PSMs in the MS Andrea results from the first replicate of the *Arabidopsis thaliana* dataset. The first three columns show the scan number, the identified peptide sequence and the delta mass (in Daltons) between the precursor and the matched peptide from the database. In the last two columns, the modification(s) determined by MS Andrea and the MSFragger localization scores are shown, respectively. The MS Andrea modifications indicate which residue is modified (single letter code and location in sequence), the modification name, the modification mass, and whether the modification is fixed or variable. The MSFragger Localization score shows the most likely location of the observed delta mass in the peptide sequence.

Here, the five top-scoring target PSMs from the MS Andrea search of the first file in the *Arabidopsis thaliana* dataset are shown. In the table, only a subset of the search result is shown; See “Data Availability” for information about where to find the full identification file. As the table shows, MS Andrea reliably identifies the peptide sequence as well as the modification(s) and on which residue(s) they are found. In contrast, MSFragger alone does not attempt to match and score potential modifications but only outputs the delta mass between the matched peptide and the precursor. Similarly, Sage will also not provide the end user with any information about the localization or type(s) of modification(s) but will also only include the difference between the precursor and matched peptide mass. MSFragger will, however, assign a localization score, which indicates the most likely position of the observed mass shift in the identified peptide sequence. If MSFragger is used in combination with PTM-Shepherd^6^, a tool which processes open search results and summarizes delta masses found by MSFragger, information about how the delta mass can be explained is also included in the output. PTM-Sheperd is developed by the same group as MSFragger and can be used within the Fragpipe Framework or as a standalone command-line interface.

For these five PSMs shown in Table 1, the localization scores that are highlighted in bold do correspond to the localization of the modifications determined by MS Andrea. For the top four spectra in Table 1, there is a general consensus between the modifications determined by MS Andrea and PTM-Shepherd. However, because PTM-Shepherd only assigns one mass shift per PSM, the two separate phoshorylations reported by MS Andrea for the spectra with scan numbers 33168 and 33407 are reported as a single pyrophosphorylation for these two spectra. For the last spectrum in the table (scan number 28757), the results differ. MS Andrea and MSFragger identify almost the same peptide, however, the peptide determined by MS Andrea has a missed tryptic cleavage and has one lysine residue more than the peptide identified by MSFragger. MS Andrea identifies one phosphorylation of threonine, while PTM-Shepherd identifies an unannotated mass shift of 208.0636 Da.

This mass roughly corresponds to the mass of a lysine residue (128.0950 Da) and one phosphorylation (79.9799 Da), indicating that this is most likely a missed identification by MSFragger in this case, despite missed cleavages being considered.

The results in Table 1 only show PSMs where a maximum of two modifications were identified by MS Andrea. However, MS Andrea is capable of identifying peptides with three and four variable modifications per peptide, which is demonstrated in Table 2.

**Table 2:**
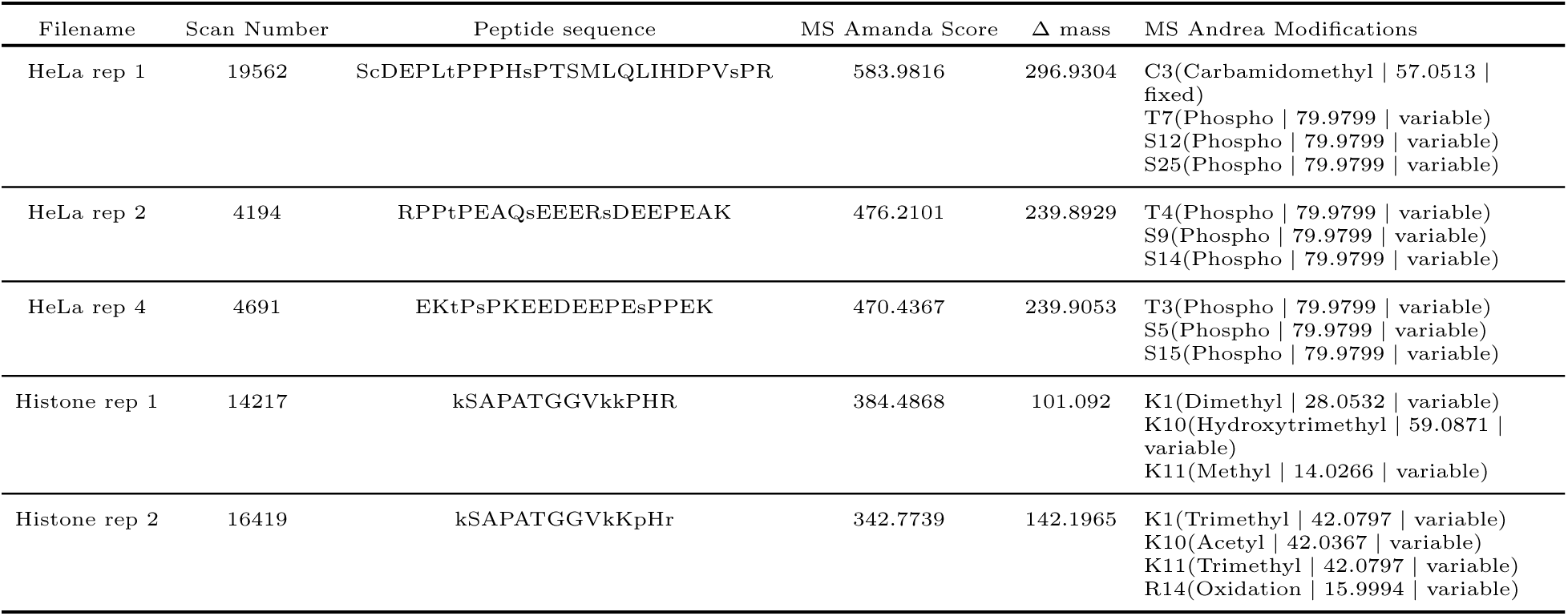
Five peptide spectrum matches (PSMs) from the HeLa and Histone dataset with more than two variable modifications identified by MS Andrea. For each PSM, the file from which it was identified, the scan number, peptide sequence, MS Amanda score, the delta mass, and the modifications determined by MS Andrea are shown.

As shown, MS Andrea can reliably identify PSMs containing up to four modifications per peptide without requiring these modifications to be predefined in the settings file, in contrast to Sage and MS Fragger. For the PSM from the first replicate of the HeLa dataset, MS Fragger identifies the same peptide sequence as MS Andrea but reports only the fixed carbamidomethyl modification, while Sage assigns a decoy peptide to this spectrum (see Supplementary Table S6 and S7). In the second replicate, all three search engines identify the same peptide sequence; however, only MS Andrea correctly localizes the modification sites. For the third PSM shown in Table 2, originating from the fourth replicate of the HeLa dataset, only MS Andrea reports a target hit for this spectrum, both Sage and MS Fragger identify decoy peptides (see Supplementary Tables S6 and S7). In addition to identifying phosphorylations, MS Andrea can also identify PTMs that are commonly found on the tails of histone proteins, as shown in the last two rows of Table 2. Two different peptidoforms of the same peptide, one with three modifications and one with four modifications, are identified in the Histone dataset. For the spectrum in the first replicate of the Histone dataset, Sage identifies a very similar, yet different, peptide as MS Andrea, with the only difference being that the second alanine is instead a serine. For that reason, the observed delta mass is also different, and the spectrum was assigned a q-value that was above the threshold of 0.01. A hit for this spectrum was not found in the output file from MSFragger with PTM-Shepherd, indicating that it also did not pass the FDR threshold. In the full MSFragger output file, the hit turned out to be a low-scoring hit to a contaminant protein. For the last spectrum shown in the table, Sage identifies a decoy hit of a contaminant, and MSFragger with PTM-Shepherd identifies a peptide from the Human Histone H4 protein rather than Histone H3. (See Supplementary Tables S6 and S7). The PSMs shown in Table 2 are examples of PSMs that were chosen to be shown here. PSMs with three and four modifications are generally observed in the MS Andrea search results. Please refer to the “Data Availability” section for information about where to find the full identification files.

These results highlight the complexity of precise modification localization in OMS. They further demonstrate that evaluating modifications at the PSM level can, in certain cases, yield high-scoring target hits and potentially increase the total number of confidently identified PSMs in a search.

### MS Andrea efficiently reduces the search space using sequence tag-based filtering

To deal with the large search space that inherently comes with OMS, MS Andrea uses a sequence tag-based filtering strategy to reduce the number of peptide candidates prior to matching and scoring. Using only the wide default mass tolerance of -500 Da and +1.005 Da, the number of peptide candidates is reduced from 2,092,395 total peptides in the database (for the FASTA file of the human proteome and based on the parameters specified in the “Search Parameters” section of the Methods) to 286,219 peptide candidates on average per spectrum for the second replicate of the HeLa dataset (see Supplementary Table S8). Using the sequence tag-based filtering in addition to filtering by mass tolerance reduces the average number of peptide candidates considered per spectrum to just 22,401. This is a reduction of 99.41% from the total number of peptides in the database and a 92.17% reduction compared to only applying the mass tolerance filtering, substantially lowering the number of considered peptide candidates and therefore the computational cost.

To evaluate the impact on final results, we ran MS Andrea without the sequence tag-based filtering step. The identification results were generally equivalent, although with a small increase in the number of identified PSMs at 1% FDR; however, at a substantially longer search time (see Supplementary Table S9). This confirms that the filtering does not sacrifice result quality while reducing the computational requirements for a search, reflected in a considerably shorter search time.

### MS Andrea Reliably Controls the False Discovery Rate

In addition to benchmarking the performance of MS Andrea against other OMS engines, we also conducted analyses to validate the FDR estimation of MS Andrea. In order to determine whether MS Andrea’s FDR control is adequate, we analyzed three files from the synthetic phosphopeptide library by Marx et al.^38^ The search results from each file were adjusted to 1% FDR using the STDA, and every PSM that did not match the peptide from the library was considered a false positive. At 1% estimated FDR, the observed true FDR was 0.64%, 0.99%, and 1.28% for the respective files, with an average value of 0.97% very close to the estimated threshold. Additionally, MS Andrea achieved a false localization rate (FLR) of 11%, 16% and 0.5%, respectively, for the three files at 1% estimated FDR. This is comparable to results obtained using a traditional closed search with MS Amanda (see Supplementary Table S10 for details). This indicates that the modification site localization of MS Andrea is on par with that of a closed database search engine, despite having the additional combinatorial complexity introduced by OMS. To further improve site localization, a dedicated modification site localization tool such as, e.g., ptmRS,^42^ could be used in the subsequent downstream analysis.

To further validate FDR control of MS Andrea, we performed an entrapment-based analysis following the framework of Wen et al. 2025,^30^ using one file from the synthetic phosphopeptide dataset. Entrapment sequences were generated with FDRBench,^30^ which adds a shuffled entrapment sequence (preserving the C-terminal residue) based on each sequence in the target FASTA file, yielding a 1:1 ratio between target and entrapment databases. The false discovery proportion (FDP) was calculated using the “combined”, “lower bound” and “sample” methods described by Wen et al. (see Supplementary Table S11). The combined-method FDP (1.05%) closely matches the 1% estimated FDR, whereas the lower bound and sample methods yield an FDP of approximately half (0.52%), which at r = 1 reflects the known underestimation of the FDP calculated using these two methods.

To test whether the localization accuracy holds on complex biological samples, we performed a search using one replicate of the HeLa dataset where phosphorylation was allowed on all 20 amino acid residues, rather than restricting phoshorylations to be found on the residues specified in the Unimod file. Extending the possible sites for phosphorylation did not compromise the localization as MS Andrea identified a phosphorylation on a false residue for only 0.88% of the PSMs at 1% estimated FDR. In general, there was a broad consensus in PSMs between the standard search and this search with all phosphorylation sites allowed (see details in Supplementary Table S12). In most of the cases where MS Andrea identified a phosphorylation on a false site, we observed that the phosphorylation was localized on a residue adjacent to a valid phosphorylation site, indicating that the false localization was most likely due to a lack of site-determining ions in the spectrum.

MS Andrea’s ability to detect PTMs that are unknown or not included in the Unimod database was tested by analysing one replicate of the synthetic phosphopeptide dataset. We repeated the search using the same search settings as the original search, but with the difference that phosphorylation was removed from the Unimod file used for the search. Despite the absence of a matching Unimod entry, MS Andrea was still able to achieve a comparable number of PSMs at 1% estimated FDR and true FDR as the standard search (See Supplementary Table S13) and identify spectra that the standard search identified as having the phosphorylation but instead as the unmodified peptidoform accompanied by a delta mass of *∼*79.979 Da, corresponding to the mass of a phosphorylation. Supplementary Figure S3 shows that the two most frequently identified delta masses for the two searches, besides 0 Da, were *∼*80 Da and *∼*98 Da, corresponding to one phosphorylation and one phosphorylation plus one oxidation, respectively. In the search in which phosphorylation was excluded, there were overall fewer PSMs for which these two delta masses were identified compared to the standard search. This is to be expected as MS Andrea directly scores and identifies modifications on the PSM level and therefore achieves a higher score when more peaks can be matched.

These analyses confirm that MS Andrea reliably controls FDR at the set threshold and that the high PSM counts reported here reflect genuine identifications.

All together, the results presented here demonstrate that MS Andrea is able to identify a higher number of PSMs (Figure 4) using STDA for FDR adjustment and comparable numbers of identifications using model-based post-processing as well-known and widely used OMS engines. Furthermore, we show that MS Andrea offers additional useful information about variable modifications and their exact locations directly on the PSM level, in contrast to Sage and MSFragger (Table 1).

## Discussion

Open modification search (OMS) extends conventional database searching by enabling the simultaneous consideration of hundreds of post-translational modifications (PTMs), which potentially can increase the identification rate of mass spectra. However, most OMS-capable search engines do not explicitly report modifications at the peptide–spectrum match (PSM) level. Instead, they report only the delta mass between the precursor and matched peptide, requiring users to manually infer the underlying modification(s) using downstream analysis tools. To address this limitation, we here presented MS Andrea, a novel OMS engine that matches and scores up to four variable modifications per peptide and reports these modifications directly at the PSM level.

We demonstrate the performance of MS Andrea on three different datasets: histones from breat cancer patients and phosphorylation-enriched peptides from HeLa cells and from *Arabidopsis thaliana*. We show that MS Andrea identifies the highest number of PSMs for the three datasets using the standard target-decoy approach (STDA) and more or comparable numbers of PSMs using model-based methods for false discovery rate (FDR) control (Figure 4 and Figure S1 and Figure S2) compared to other OMS engines, namely MS Fragger and Sage. On the Histone dataset, MS Andrea, Sage, and MSFragger agreed on a core set of PSMs while each also contributed a notable number of unique identifications, with MS Andrea identifying the most (Figure 5). Although the overlap between engines was modest, closer inspection showed that some of the discrepancy between the search engines arises from differences in target–decoy classification rather than from entirely distinct peptides being assigned to the spectra, indicating that the three search engines agree more closely at the level of PSMs than the raw overlap suggests.

Where MS Andrea stands out compared to other OMS engines is its ability to score and match variable modifications in the PSM level. We show this here for the five top-scoring PSMs from the first replicate of the *Arabidopsis thaliana* dataset (Table 1). While MS Andrea and MSFragger both largely identify the same peptide sequence and delta mass, MS Andrea will decipher this delta mass and assign modification(s) that correspond to this mass to the peptide. This is in contrast to MSFragger, which alone only provides the user with a localization score for the observed delta mass and depends on additional tools such as PTM-Shepherd to achieve similar results as MSAndrea. Even when PTM-Shepherd is employed in combination with MSFragger, it will still explain the observed mass shift as one modification, rather than a combination of modifications, leading to the assignment of pyrophosphorylations rather than two separate phosphorylations as assigned by MS Andrea for the *Arabidopsis thaliana*. (Table 1) Additionally, we show that MS Andrea can confidently identify peptides with up to four variable modifications of various types, including phosphorylations and modifications commonly found on the tails of histone proteins (Table 2). Additionally, we demonstrate that the sequence tag-based filtering of MS Andrea efficiently reduces the search space and therefore reduces computational demands without compromising the quality of the results. Lastly, the validity of these identifications is supported by empirical FDR validation and entrapment analysis on the synthetic library of phosphorylated peptides, confirming that MS Andrea can reliably control the FDR at this threshold.

Currently, MS Andrea is available as a command line tool for Windows, Linux and macOS. However, we are planning to implement a node for MS Andrea in the Proteome Discoverer Software (Thermo Fischer Scientific) in the future.

## Data Availability

MS Andrea is implemented in C# and is currently available as a command-line interface for Windows, Linux and macOS and can be downloaded for free at https://github.com/hgb-bin-proteomics/MSAndrea. As input, MS Andrea takes MS2 spectra in either .mgf or .mzml format, and the output is provided as .csv.

Result files from all three search engines, pin and pout files based on results from MS Andrea and MS Fragger and the analyzed .mzml files are available on Zenodo at https://doi.org/10.5281/zenodo.21902795.

Datasets used for benchmarking MS Andrea were made available by the original authors via the PRIDE repository^32^ (https://www.ebi.ac.uk/pride/), using the following project identifiers:

- HeLa dataset, Bekker-Jensen et al. 2020^34^ : PXD014525
- Arabidopsis dataset, Mergner et al. 2020^36^ : PXD013868
- Histone dataset, Noberini et al. 2016^37^ : PXD002669
- Synthetic Phosphorylated Peptide Library, Marx et al. 2013^38^ : PXD000138

## Author Contributions

Louise Marie Buur: conceptualization, methodology, software, visualization, writing - original draft

Stephan Winkler: writing - review and editing, supervision

Viktoria Dorfer: conceptualization, methodology, software, writing - review and editing, supervision, funding acquisition

## Acknowledgement

L.M.B. acknowledges the MSCA-ITN-2020 PROTrEIN project that has received funding from the European Union’s Horizon 2020 programme under the Marie Sk-lodowska-Curie grant agreement N° 956148.

We thank Micha J. Birklbauer, Sebastian Dorl, Susanne Schaller and Marina Strobl from the Bioinformatics Research Group and Manuel Matzinger from the Proteomics Technology Hub, IMP, Vienna for their fruitful discussions and valuable input to the project and the manuscript.

## Supporting Information Available

Supporting information is available online at [INSERT LINK]. The supporting information (PDF) includes additional experimental details and results, as listed below:

- Supplementary Element S1: Description of filters applied to create the modification dictionary by MS Andrea
- Supplementary Element S2: Full list of analyzed files from the HeLa dataset
- Supplementary Element S3: Full list of analyzed files from the *Arabidopsis thaliana* dataset
- Supplementary Element S4: Full list of analyzed files from the Histone dataset
- Supplementary Element S5: Full list of analyzed files from the Synthetic Phosphorylated Peptide Library
- Supplementary Table S1: Full list of added auxiliary peaks before sequence tag extraction in MS Andrea
- Supplementary Table S2: List of default modifications considered by MS Andrea
- Supplementary Table S3: Average number of identified PSMs and standard deviations for the Histone dataset
- Supplementary Table S4: Average number of identified PSMs and standard deviations for the *Arabidopsis thaliana* dataset
- Supplementary Table S5: Average number of identified PSMs and standard deviations for the HeLa dataset.
- Supplementary Table S6: PSMs from Sage for the same five PSMs shown in Table 2 of the main manuscript
- Supplementary Table S7: PSMs from MSFragger for the same five PSMs shown in Table 2 of the main manuscript
- Supplementary Table S8: Metrics showing efficiency of the sequence tag-based filtering strategy
- Supplementary Table S9: Number of PSMs with and without sequence tag-based filtering
- Supplementary Table S10: False Discovery-and Localization Rate for MS Amanda search
- Supplementary Table S11: False Discovery Proportions for Entrapment Analysis
- Supplementary Table S12: Overlap in PSMs at 1% FDR between Standard MS Andrea and All Phopsho MS Andrea search
- Supplementary Table S13: Number of PSMs and true FDR for Standard MS Andrea and No Phospho MS Andrea search
- Supplementary Figure S1: Average number of PSMs at 1% FDR for the *Arabidopsis thaliana* dataset.
- Supplementary Figure S2: Average number of PSMs at 1% FDR for the HeLa dataset.
- Supplementary Figure S3: Delta masses identified by MS Andrea for one replicate of the synthetic phosphopeptide library

## TOC Graphic

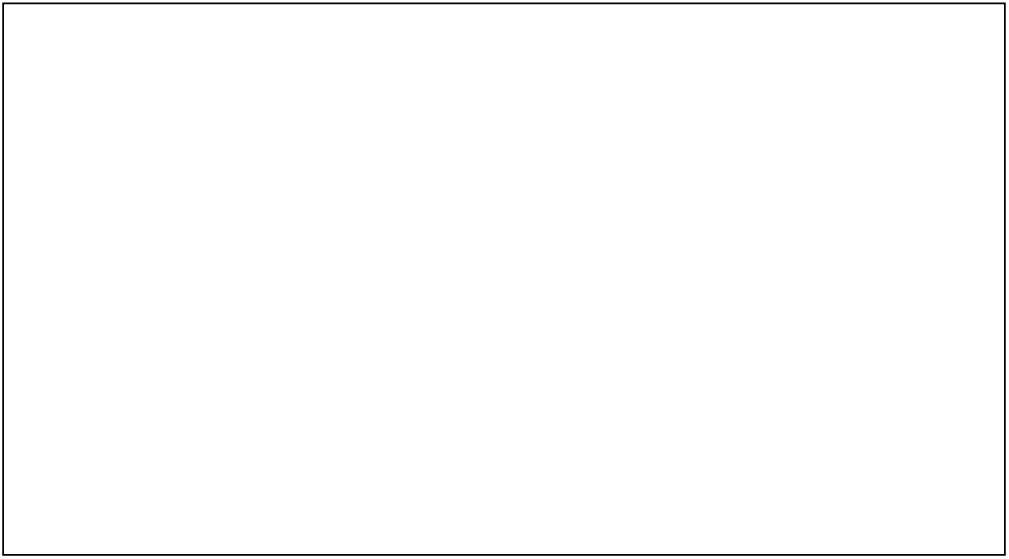

