## Supporting Information for "Beyond Delta Masses: MS Andrea Directly Resolves Combinatorial Peptide Modifications in Open Searches"

Louise Marie Buur[1,2], Stephan Winkler[1,2] and Viktoria Dorfer*[,1]

[1] Bioinformatics Research Group, University of Applied Sciences Upper Austria,

4232 Hagenberg, Austria

[2] Institute for Symbolic Artificial Intelligence, Johannes Kepler University Linz, 4040 Linz, Austria

*

#### Table of Contents

Supplementary Element S1: Description of filters applied to create the modification dictionary by MS Andrea .............................. S2

Supplementary Element S2: Full list of analyzed files from the HeLa dataset.................................................................................. S3

Supplementary Element S3: Full list of analyzed files from the *Arabidopsis thaliana* dataset......................................................... S3

Supplementary Element S4: Full list of analyzed files from the Histone dataset............................................................................ S3

Supplementary Element S5: Full list of analyzed files from the Synthetic Phosphorylated Peptide Library..................................... S3

Supplementary Table S1: Full list of added auxiliary peaks before sequence tag extraction in MS Andrea ..................................... S4

Supplementary Table S2: List of default modifications considered by MS Andrea .......................................................................... S5

Supplementary Table S3: Average number of identified PSMs and standard deviations for the Histone dataset.............................. S6

Supplementary Table S4: Average number of identified PSMs and standard deviations for the *Arabidopsis thaliana* dataset............ S6

Supplementary Table S5: Average number of identified PSMs and standard deviations for the HeLa dataset.................................. S6

Supplementary Table S6: PSMs from Sage for the same five PSMs shown in Table 2 of the main manuscript.................................. S7

Supplementary Table S7: PSMs from MSFragger for the same five PSMs shown in Table 2 of the main manuscript .......................... S7

Supplementary Table S8: Metrics showing efficiency of the sequence tag-based filtering strategy................................................. S8

Supplementary Table S9: Number of PSMs with and without sequence tag-based filtering .......................................................... S9

Supplementary Table S10: False Discovery- and Localization Rate for MS Amanda search .......................................................... S9

Supplementary Table S11: False Discovery Proportions for Entrapment Analysis.......................................................................... S9

Supplementary Table S12: Overlap in PSMs at 1% FDR between Standard MS Andrea and "All Phopsho" MS Andrea search ..........S10

Supplementary Table S13: Number of PSMs and true FDR for Standard MS Andrea and "No Phospho" MS Andrea search..............S10

Supplementary Figure S1: Average number of PSMs at 1% FDR for the *Arabidopsis thaliana* dataset ............................................S11

Supplementary Figure S2: Average number of PSMs at 1% FDR for the HeLa dataset ...................................................................S12

Supplementary Figure S3: Delta masses identified by MS Andrea for one replicate of the synthetic phosphopeptide library ...........S13

References ...................................................................................................................................................S14

#### Supplementary Element S1: Description of filters applied to create the modification dictionary by MS Andrea

Based on the Unimod .xml file, MS Andrea constructs a dictionary of possible combinations of up to four modifications. As of June 2026, the Unimod database contains 1358 modifications. Considering the full set of C(1358, 4) ≈ 1.41 × $10^{11}$ combinations is not computationally feasible within a reasonable amount of time. Therefore, MS Andrea applies four filters before the modification dictionary is built. The four filters are:

1. Substitutions (modifications with “->” in the title field) are excluded as these represent amino acid variants rather than PTMs
2. Only modifications with at least one residue classified as “Post-translational” are included. This restricts the dictionary to biologically occurring modifications and excludes isotopic labels and chemical derivatives. Even though only one site for a given modification is classified as “Post-translational” all residues are treated as potential sites even though that site may have a different classification
3. During construction of the dictionary, only combinations whose combined mass falls within the open-search precursor mass window (bounded by the values specified for ms1_tol_open_search_lower and ms1_tol_open_search_upper) are generated. This means that combinations that do not correspond to any observable delta mass are never enumerated
4. Combinations of modification types with mutually cancelling masses are excluded from the dictionary, e.g., any combination containing both “oxidation” and “deoxy” (per the Unimod.xml naming). Although such complementary modifications can co-occur on a peptide in principle, in practice we found them to be predominantly false positives (e.g., an oxidation on one residue paired with a deoxidation on the adjacent residue), and we therefore decided to exclude such cases

Applying these filters reduce the number of modifications from 1358 to 133 commonly observed, relevant modifications, and the number of four-modification combinations from ~1.41 × $10^{11}$ to C(133, 4) = 12,457,445. This number is then further reduced to 1,902,572 by applying the default mass tolerance settings (-500 Da and + 1.005 Da) and removing combinations with masses that cancel out each other. A full list of all 133 modifications considered by MS Andrea by default is shown in Supplementary Table S2.

#### Supplementary Element S2: Full list of analyzed files from the HeLa dataset

- 20171001_QE3_nLC7_DBJ_SA_LFQphos_LCMS_Rep_01.raw
- 20171001_QE3_nLC7_DBJ_SA_LFQphos_LCMS_Rep_02.raw
- 20171001_QE3_nLC7_DBJ_SA_LFQphos_LCMS_Rep_03.raw
- 20171001_QE3_nLC7_DBJ_SA_LFQphos_LCMS_Rep_04.raw
- 20171001_QE3_nLC7_DBJ_SA_LFQphos_LCMS_Rep_05.raw
- 20171001_QE3_nLC7_DBJ_SA_LFQphos_LCMS_Rep_06.raw

The data was originally published by Bekker-Jensen et al. in 2020 [1] . The full dataset is available via the PRIDE repository (https://www.ebi.ac.uk/pride/) under identifier PXD014525.

#### Supplementary Element S3: Full list of analyzed files from the *Arabidopsis thaliana* dataset

- 02131_A01_P022360_I00_U01_R1.raw
- 02131_B01_P022360_I00_U02_R1.raw
- 02131_C01_P022360_I00_U03_R1.raw
- 02131_D01_P022360_I00_U04_R1.raw
- 02131_E01_P022361_I00_U01_R1.raw
- 02131_F01_P022361_I00_U02_R1.raw
- 02131_G01_P022361_I00_U03_R1.raw
- 02131_H01_P022361_I00_U04_R1.raw

The data was originally published by Mergner et al. 2020 [2] . The full dataset is available via the PRIDE repository under identifier PXD013868.

#### Supplementary Element S4: Full list of analyzed files from the Histone dataset

- F140814 Sample3 Frozen H3 01.raw
- F140814 Sample3 Frozen H3 02.raw
- F140814 Sample3 Frozen H3 03.raw

The data was originally published by Noberini et al. 2016 [3] . The full dataset is available via the PRIDE repository under identifier PXD002669.

#### Supplementary Element S5: Full list of analyzed files from the Synthetic Phosphorylated Peptide Library

- 18.raw
- 21.raw
- 50.raw

The data was originally published by Marx et al. 2013 [4] . The full dataset is available via the PRIDE repository under identifier PXD000138.

#### Supplementary Table S1: Full list of added auxiliary peaks before sequence tag extraction in MS Andrea

*Table S1. Auxiliary peaks acting as the start and end point of the ions series.*

| Ion type | Added peaks |
|---|---|
| b | $b_0: \ m_{proton}$<br>$b_{end}: \ (m_{precursor} + m_{proton}) - m_{H_2O}$ |
| y | $y_0: \ m_{proton} + m_{H_2O}$<br>$y_{end}: \ m_{precursor} + m_{proton}$ |
| c | $c_0: \ m_{proton} + NH_3$<br>$c_{end}: \ (m_{precursor} + m_{proton}) - m_{H_2O} + m_{NH_3}$ |
| z | $z + 1_0: \ m_{proton} + m_{H_2O} - NH$<br>$z + 2_0: \ m_{proton} + m_{proton} + m_{H_2O} - NH$<br>$z_{end}: \ (m_{precursor} + m_{proton}) - m_{NH}$ |

### Supplementary Table S2: List of default modifications considered by MS Andrea

*Table S2. All modifications considered by default by MS Andrea when creating the dictionary of combined modifications. Each of these modifications have at least one residue which is categorized as "Post-translational".*

| | | |
|---|---|---|
| (-)-dehydroepigallocatechin | Glycerophospho | phycocyanobilin |
| (-)-epigallocatechin-3-gallate | glycerylphosphorylethanolamine | phycoerythrobilin |
| 13-dioxo-7-heptadecenoic ester | glyoxal-derived AGE | phosphoribosyl dephospho-coenzyme A |
| 2-amino-3-oxo-butanoic_acid | heme | Phosphorylation |
| 2-hydroxyisobutyrylation | hydrogenase diiron subcluster | phytochromobilin |
| 3,4-didehydroretinylidene | hydroxycinnamyl | Pyridoxal phosphate |
| 3-phosphoglyceryl | hydroxyfarnesyl | pyrophosphorylation of Ser/Thr |
| 4-didehydroretinylidene | hydroxyheme | quinone |
| 4-hydroxynonenal (HNE) | hydroxymethyl | Reaction with dimethylarsinous (AsIII) acid |
| 4-methyl-delta-1-pyrroline-5-carboxyl | hypusine | reduction |
| 5-glutamyl serotonin | L-selenocysteinyl molybdenum bis(molybdopterin guanine dinucleotide) | S-(4a-FMN) |
| 5-hydroxy-N6,N6,N6-trimethyl | LeudimethylArgGlyGly | S-diphytanylglycerol diether |
| Acetylation | LeumethylArgGlyGly | S-guanylation-2 |
| Acetyldeoxyhypusine | Levuglandinyl-lysine anhydrolactam adduct | S-homocysteinylation |
| Acetylhypusine | Levuglandinyl-lysine anhyropyrrole adduct | S-nitrosylation |
| Addition of DEDGFLYMVYASQETFG | Levuglandinyl-lysine pyrrole adduct | selenyl |
| addition of GGE | Levuglandinyl - arginine hydroxylactam adduct | Succinic anhydride labeling reagent light form (N-term & K) |
| adduction of aflatoxin B1 Dialdehyde to lysine | Levuglandinyl - arginine lactam adduct | SulfurDioxide |
| amidino | Levuglandinyl - lysine hydroxylactam adduct | SUMOylation by Giardia lamblia |
| Beta-methylthiolation | Levuglandinyl - lysine lactam adduct | tetraglutamyl |
| Biotinylation | lipid | tetraiodo |
| bromination | Lipoyl | tri-Methylation |
| Butyryl | Loss of ammonia | triglutamyl |
| Carbamylation | Loss of ammonia (15N) | triiodo |
| carboxyethyl | maleimide-3-saccharide | Ubiquitin D (FAT10) leaving after chymotrypsin digestion Cys-Ile-Gly-Gly |
| Carboxylation | maleimide-5-saccharide | Ubiquitin D (FAT10) leaving after trypsin digestion |
| cis-14-hydroxy-10 | Malonylation | Ubiquitination 2H4 lysine |
| copper sulfido molybdopterin cytosine dinuncleotide | Methylation | ubiquitinylation residue |
| Crotonylation | Modification by hydroxylated mechloroethamine (HN-2) | uridine phosphodiester |
| cyano | Modification by hydroxylated tris-(2-chloroethyl)amine (HN-3) | |
| Cysteine modified Coenzyme A | molybdenum bis(molybdopterin guanine dinucleotide) | |
| Cytidine monophosphate | molybdopterin | |
| Deamidation followed by a methylation | monoglutamyl | |
| Dehydration | Myristoylation | |
| Deoxyhypusine | N-acyl diglyceride cysteine | |
| di-Methylation | N-Homocysteine thiolactone | |
| diacylglycerol | N-pyruvic acid 2-iminyl | |
| diglutamyl | Nitroalkylation by Nitro Linoleic Acid | |
| dihydroxy | Nitroalkylation by Nitro Oleic Acid | |
| Dimethylation of proline residue | O-Sulfonation | |
| Diphthamide | O3-(riboflavin phosphoryl) | |
| dipyrrolylmethanemethyl | octanoyl | |
| DMPO spin-trap nitrone adduct | Oxidation or Hydroxylation | |
| Enzymatic glycine removal leaving an amidated C-terminus | palmitoleyl | |
| Farnesylation | Palmitoylation | |
| Flavin adenine dinucleotide | persulfide | |
| flavin mononucleotide | phosphate-ribosylation | |
| Formation of five membered aromatic heterocycle | phospho-guanosine | |
| Geranyl-geranyl | Phosphogluconoylation | |
| glutathione disulfide | Phosphopantetheine | |

#### Supplementary Table S3: Average number of identified PSMs and standard deviations for the Histone dataset

*Table S3. Average number of identified PSMs and standard deviation for the Histone dataset, which are shown in the main manuscript in Figure 4. Shown is the search engine name, false discovery rate (FDR) method, either standard target-decoy approach (STDA) or machine learning based using Percolator or the q-values intrinsically determined by Sage.*

| Search engine | FDR method | Average number of PSMs | Standard deviation |
|---|---|---|---|
| MS Andrea | STDA | 539 | 6 |
| MS Andrea | Percolator | 1000 | 84 |
| MSFragger | STDA | 285 | 69 |
| MSFragger | PeptideProphet | 418 | 12 |
| Sage | STDA | 24 | 5 |
| Sage | Sage q-value | 677 | 82 |

#### Supplementary Table S4: Average number of identified PSMs and standard deviations for the *Arabidopsis thaliana* dataset

*Table S4. Average number of identified PSMs and standard deviation for the Arabidopsis thaliana dataset, which are shown here in the Supplementary material in Figure S1. Shown is the search engine name, false discovery rate (FDR) method, either standard target-decoy approach (STDA) or machine learning based using Percolator or the q-values intrinsically determined by Sage.*

| Search engine | FDR method | Average number of PSMs | Standard deviation |
|---|---|---|---|
| MS Andrea | STDA | 6968 | 1558 |
| MS Andrea | Percolator | 9795 | 1807 |
| MSFragger | STDA | 6580 | 1628 |
| MSFragger | PeptideProphet | 10781 | 1598 |
| Sage | STDA | 2760 | 826 |
| Sage | Sage q-value | 10131 | 1923 |

#### Supplementary Table S5: Average number of identified PSMs and standard deviations for the HeLa dataset

*Table S5. Average number of identified PSMs and standard deviation for the HeLa dataset, which are shown here in the Supplementary material in Figure S2. Shown is the search engine name, false discovery rate (FDR) method, either standard target-decoy approach (STDA) or machine learning based using Percolator or the q-values intrinsically determined by Sage.*

| Search engine | FDR method | Average number of PSMs | Standard deviation |
|---|---|---|---|
| MS Andrea | STDA | 6110 | 102 |
| MS Andrea | Percolator | 8305 | 311 |
| MSFragger | STDA | 5219 | 128 |
| MSFragger | PeptideProphet | 9590 | 151 |
| Sage | STDA | 1547 | 102 |
| Sage | Sage q-value | 6320 | 66 |

#### Supplementary Table S6: PSMs from Sage for the same five PSMs shown in Table 2 of the main manuscript

*Table S6. Five peptide spectrum matches from Sage for the same five spectra shown in the main manuscript in Table 2. Shown is the filename, scan number, identified peptide sequence, delta mass and the protein in which the peptide is found.*

| Filename | Scan Number | Peptide sequence | Δ mass | (target/decoy/contaminant) |
|---|---|---|---|---|
| HeLa rep 1 | 19562 | IGGGNVPGRSGRPLTEEQLLQQQQQQLHR | 23.67 | Decoy |
| HeLa rep 2 | 4194 | RPPTPEAQSEEERSDEEPEAK | 239.893 | Target |
| HeLa rep 4 | 4691 | AIPSTSIRAAPSVSRVPSPTPR | 174.652 | Decoy |
| Histone rep 1 | 14217 | KSAPSTGGVKKPHR | 85.097 | Target |
| Histone rep 2 | 16419 | TSGTKPAVASCSSFR | 77.221 | Decoy, contaminant |

#### Supplementary Table S7: PSMs from MSFragger for the same five PSMs shown in Table 2 of the main manuscript

*Table S7. Five peptide spectrum matches from MSFragger for the same five spectra shown in the main manuscript in Table 2. Shown is the filename, scan number, identified peptide sequence, delta mass and the protein in which the peptide is found.*

| Filename | Scan Number | Peptide sequence | Δ mass | (target/decoy/contaminant) |
|---|---|---|---|---|
| HeLa rep 1 | 19562 | SCDEPLTPPPHSPTSMLQLIHDPVSPR | 239.9087 | Target |
| HeLa rep 2 | 4194 | RPPTPEAQSEEERSDEEPEAK | 239.8931 | Target |
| HeLa rep 4 | 4691 | KEPPSEPEEDEEKPSPTKEK | 174.6520 | Decoy |
| Histone rep 1 | 14217 | IAVGGFR | 107.0472 | Contaminant |
| Histone rep 2 | 16419 | GKGGKGLGKGGAKRHR | 12.0229 | Target |

**MS Fragger Localization score, HeLa rep 1, scan number 19562:**

S(6.60)C(6.60)D(9.56)E(11.11)P(12.74)L(14.18)T(15.84)P(15.84)P(15.84)P(15.84)H(15.84)S(15.84)P(15.84)T(15.84)
S(15.84)M(15.84)L(15.84)Q(15.84)L(15.84)I(15.84)H(15.84)D(15.84)P(14.67)V(14.67)S(14.67)P(11.87)R(9.24)

**MS Fragger Localization score, HeLa rep 2, scan number 4194:**

R(15.78)P(17.49)P(18.97)T(18.97)P(20.09)E(20.09)A(20.09)Q(20.09)S(20.09)E(20.09)E(20.09)E(20.09)R(20.09)S(20.09)D(18.12)E(16.27)E(16.24)P(14.09)E(12.67)A(10.33)K(7.25)

**MS Fragger Localization score, HeLa rep 4, scan number 4691:**

K(0.66)E(2.27)P(3.95)P(3.95)S(3.95)E(5.86)P(5.86)E(7.86)E(9.86)D(11.77)E(13.75)E(15.87)K(18.11)P(18.11)S(20.47)P(20.47)T(20.47)K(20.47)E(20.47) K(19.81)

**MS Fragger Localization score, Histone rep 1, scan number 14217:**

I(2.88)A(2.88)V(7.01)G(5.36)G(5.36)F(5.36)R(5.14)

**MS Fragger Localization score, Histone rep 2, scan number 16419:**

G(4.51)K(4.51)G(5.31)G(5.31)K(5.31)G(5.31)L(0.52)G(5.91)K(8.82)G(10.35)G(10.35)A(10.35)K(10.35)R(10.35)H(10.35)R(9.17)

#### Supplementary Table S8: Metrics showing efficiency of the sequence tag-based filtering strategy

To investigate the impact on the sequence tag-based filtering strategy, we slightly modified MS Andrea to export the number of peptide candidates for each spectrum using three different filtering strategies (mass tolerance only, sequence tags only and mass tolerance in combination with sequence tags). We analyzed the second replicate of the HeLa dataset using the parameters specified under "Search Parameters" of the Methods in the main manuscript. We then calculated the mean and median number of peptide candidates as well as the standard deviation and percent reduction in peptides based on the total number.

*Table S8. Summary of peptide candidate reduction across three filtering strategies applied to the peptide database based on the parameters described in the 'Search Parameters' section of the main manuscript. The data is shown for spectra from the second replicate of the HeLa dataset. For each strategy, mass tolerance (mass tol.) only, sequence tags (seq. tags) only, and combined mass tol. and seq. tags, the mean, median, and standard deviation (Std. dev.) of remaining candidates per spectrum are reported, along with the percentage reduction of the search space relative to the total number of peptides in the database.*

| Total number of peptides in database | | Filtering using only mass tol. | Filtering using only seq. tags | Filtering using both seq. tags and mass tol. |
|---|---|---|---|---|
| 2,092,395 | Mean | 286,219 | 168,975 | 22,401 |
| | Median | 307,230 | 116,253 | 12,408 |
| | Std. dev | 121,092 | 168,518 | 26,552 |
| | % reduced from total peptides (based on mean) | 83.32 % | 91.92 % | 99.41 % |
| | % reduced from filtering using only mass tol. (based on mean) | x | 40.96 % | 92.17 % |

#### Supplementary Table S9: Number of PSMs with and without sequence tag-based filtering

*Table S9. Number of peptide spectrum matches (PSMs) at 1% false discovery rate (FDR) using the standard target-decoy approach (STDA) and percolator for FDR estimation along with search time for the second replicate of the HeLa dataset. The file was analyzed with three different versions of MSAndrea. Standard MS Andrea, MS Andrea where the sequence tag-based filtering was disabled and lastly MS Andrea where only sequence tags of two and three residues were used for filtering. The parameters for each search here the same and can be found in the "Search Parameters" section of the Methods in the main manuscript.*

| | **Number of PSMs at 1% FDR (STDA)** | **Number of PSMs at 1% FDR (Percolator)** | **Search time** |
|---|---|---|---|
| Standard MS Andrea | 6225 | 8697 | 164 minutes |
| MS Andrea without sequence tag-based filtering | 6375 | 9143 | 341 minutes |

#### Supplementary Table S10: False Discovery- and Localization Rate for MS Amanda search

*Table 10. False discovery rate (FDR) calculated using the standard target-decoy approach and false localization rate (FLR) for the MS Amanda search of the three files (18.raw, 21.raw and 50.raw) from the synthetic phosphopeptide library. The MS Amanda search was conducted using the same search parameters as for MS Andrea, with the only difference being that phosphorylation of S, T and Y was set as a variable modification.*

| | **18.raw** | **21.raw** | **50.raw** |
|---|---|---|---|
| FDR | 1.91 % | 4.1 % | 2.5 % |
| FLR | 12 % | 13 % | 0.5 % |

#### Supplementary Table S11: False Discovery Proportions for Entrapment Analysis

We conducted an entrapment analysis of the "18.raw" file from the synthetic phosphopeptide dataset by Marx et al. to further confirm the FDR control of MS Andrea. We followed the framework published by Wen et al. 2025 [5] and calculated the false discovery proportion (FDP) using the three following methods described by the authors: (1) "combined" method: $FDP = \left(\frac{E*(1+1/r)}{T+E}\right)$, (2) "lower bound" method: $FDP = \left(\frac{E}{T+E}\right)$ and (3) "sample" method: $FDP = \left(\frac{E*(1/r)}{T}\right)$. Entrapment sequences were added to the FASTA file using FDRBench v. 1.1.1, released by Wen et al. in connection with their publication. FDRBench added a shuffled sequence of each entry in the FASTA file (preserving the C-term residues), resulting in a ratio of 1:1 between the target database and the entrapment database. Shown in Table S11 is the FDP using the three different methods at the estimated FDR threshold of 1%.

*Table S11. Estimated false discovery rate (FDR) and false discovery proportion (FDP) for the entrapment analysis of the "18.raw" files calculated using the "combined", "lower bound" and "sample" methods as mentioned by Wen et al. 2025 [5].*

| **Estimated FDR** | **Combined FDP** | **Lower bound FDP** | **Sample FDP** |
|---|---|---|---|
| 1,00 % | 1,0477 % | 0,5239 % | 0,5226 % |

#### Supplementary Table S12: Overlap in PSMs at 1% FDR between Standard MS Andrea and All Phopsho MS Andrea search

We tested the localization accuracy of MS Andrea on the second replicate of the HeLa dataset by performing a search with MS Andrea where phosphorylation was allowed on all 20 amino acid residues (All Phospho), rather than restricting phosphorylations to be found on the residues specified in the Unimod file (Standard). The number of PSMs at 1% estimated FDR for the Standard and All Phospho searches are shown in Table S12.

*Table S12.Number of PSMs at 1% estimated FDR for the Standard MS Andrea and "All Phospho" MS Andrea search for the second replicate of the HeLa dataset. FDR was estimated using the standard target-decoy approach.*

| Number of PSMs for which there is an overlap between the two searches | Number of PSMs unique to Standard MS Andrea | Number of PSMs unique to All Phosho MS Andrea |
|---|---|---|
| 6130 | 156 | 143 |

Out of the 156 and 143 PSMs that are unique to each of the searches, 55 of them are spectra for which the "All Phospho" search found a phosphorylation on a false residue (which corresponds 0.88% of spectra at 1% estimated FDR), which is why there is no overlap for these spectra between the searches. For the remaining spectra (101 and 88, respectively) there was no hit for these spectra in the other search within the threshold at 1% FDR. Further investigation showed that the PSMs for these spectra were either decoy hits or target hits with a score below the threshold.

#### Supplementary Table S13: Number of PSMs and true FDR for Standard MS Andrea and No Phospho MS Andrea search

To test the ability of MS Andrea to identify unknown PTMs or PTMs not present in the Unimod file, we analysed the "18.faw" file of the synthetic phosphorylated peptide library using with MS Andrea where phosphorylation was removed from the Unimod file used for the search (No Phospho MS Andrea). Shown in Table S13 are the number of PSMs at 1 % estimated FDR using the standard target-decoy approach and the true FDR for the Standard and the No Phospho MS Andrea.

*Table S13. Number of PSMs at 1 % estimated FDR and true FDR for the "18.raw" file of the synthetic phosphopeptide dataset for the Standard MS Andrea search and the No Phospho Search where phosphorylation was excluded from the Unimod file.*

| | Standard MS Andrea | No Phospho MS Andrea |
|---|---|---|
| Number of PSMs at 1% estimated FDR | 5188 | 5178 |
| True FDR | 0.636 % | 0.637 % |

The delta masses identified by the two respective searches at 1% estimated FDR are shown in Supplementary Figure S3.

#### Supplementary Figure S1: Average number of PSMs at 1% FDR for the *Arabidopsis thaliana* dataset

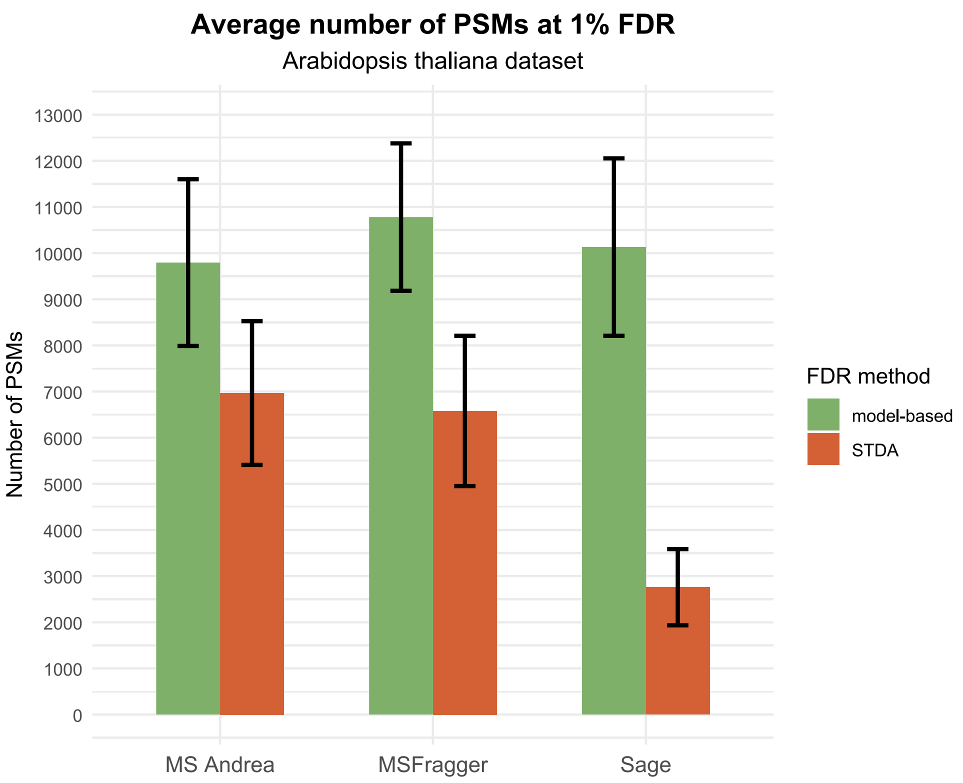


*Figure S1. Average number of PSMs determined by MS Andrea, MSFragger and Sage for the eight replicates of the Arabidopsis Thaliana dataset. Green bars show the average number of PSMs for the search engine with machine learning (ML)-based post-processing. For MS Andrea post-processing was done with Percolator, and with PeptideProphet for MSFragger. For Sage results, the q-value determined on spectrum level by Sage was used. Orange bars show the average number of PSMs at 1% false discovery rate (FDR) using the standard target-decoy approach (STDA). Error bars show standard deviation among the replicates.*

#### Supplementary Figure S2: Average number of PSMs at 1% FDR for the HeLa dataset

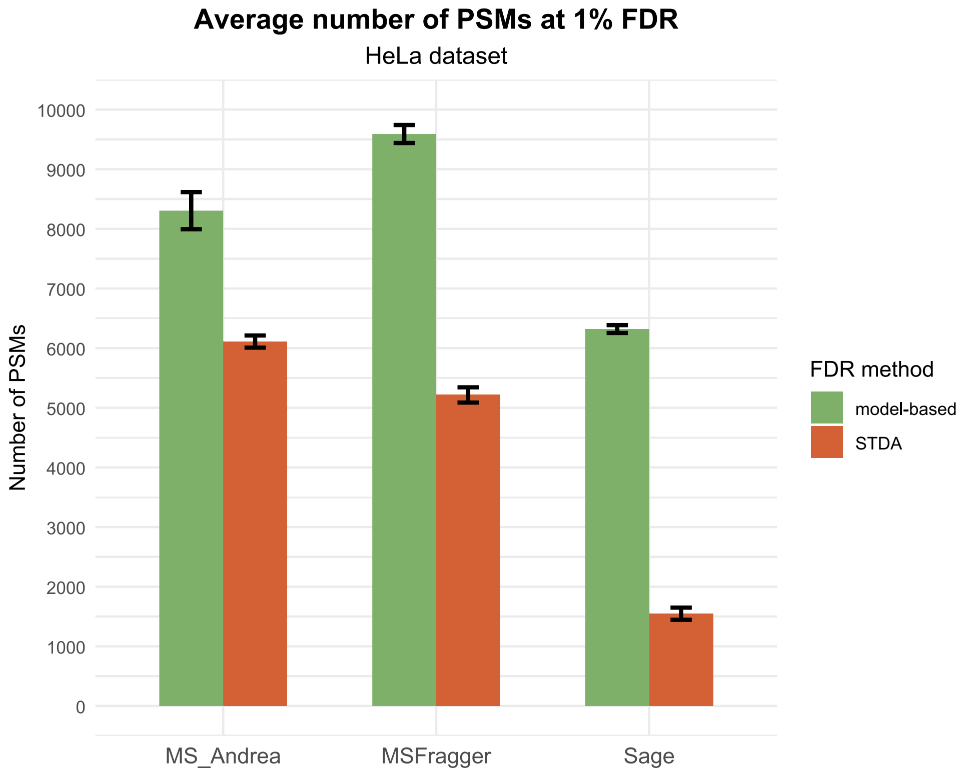


*Figure S2. Average number of PSMs determined by MS Andrea, MSFragger and Sage for the six replicates of the HeLa dataset. Green bars show the average number of PSMs for the search engine with machine learning (ML)-based post-processing. For MS Andrea post-processing was done with Percolator, and with PeptideProphet for MSFragger. For Sage results, the q-value determined on spectrum level by Sage was used. Orange bars show the average number of PSMs at 1% false discovery rate (FDR) using the standard target-decoy approach (STDA). Error bars show standard deviation among the replicates.*

#### Supplementary Figure S3: Delta masses identified by MS Andrea for one replicate of the synthetic phosphopeptide library

To test the ability of MS Andrea to identify unknown PTMs or PTMs not present in the Unimod file, we analysed the "18.faw" file of the synthetic phosphorylated peptide library using with MS Andrea where phosphorylation was removed from the Unimod file used for the search (No Phospho MS Andrea). Shown in Figure S3 are the delta masses identified by the Standard MS Andrea search and the No Phospho MS Andrea search at 1% estimated FDR.

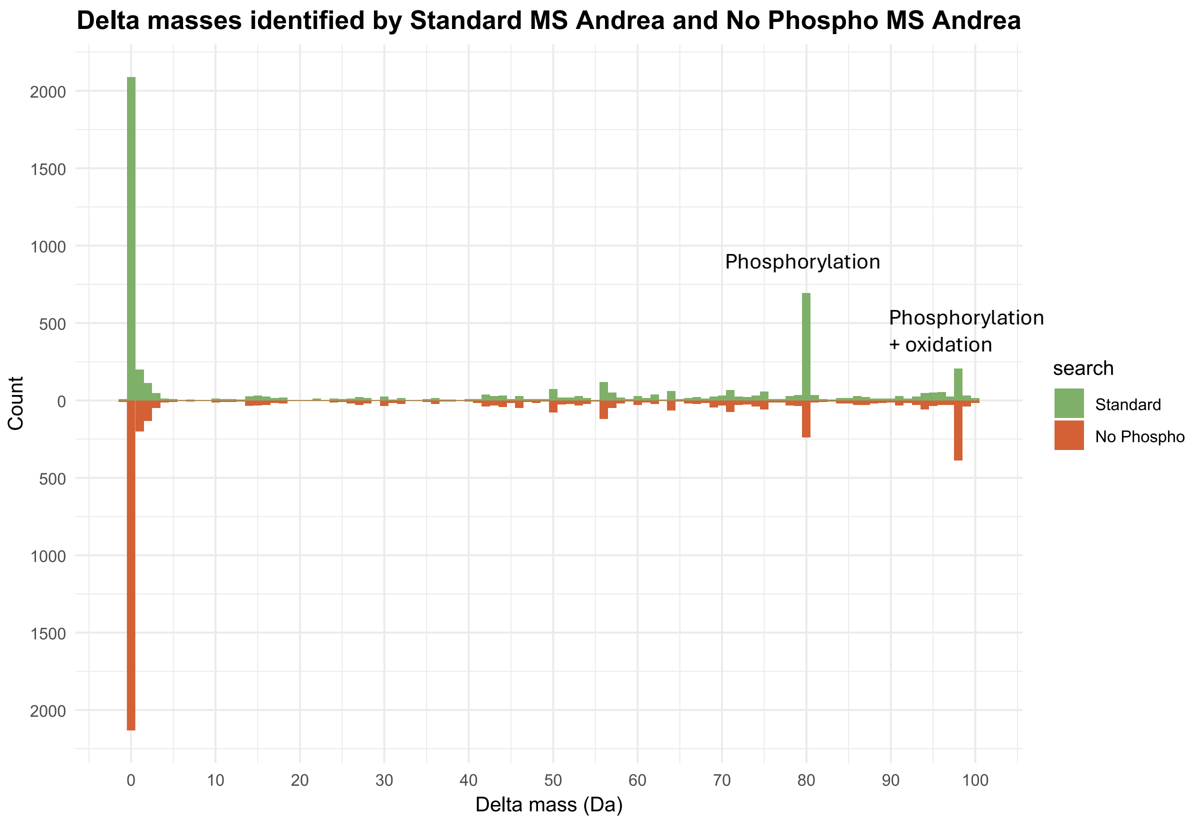


*Figure S3. Delta masses (in Da) identified by Standard MS Andrea (green) and No Phopsho MS Andrea (orange), where phosphorylation was excluded from the Unimod file used in the search, for the "18.faw" file from the synthetic phosphopeptide dataset. Indicated on the plot are the masses that correspond to one phosphorylation (~80 Da) and one phosphorylation plus one oxidation (~98 Da).*

The number of PSMs identified and 1% FDR and the true FDR for the two respective searches are shown in Supplementary Table S13.
